# Astrocytic mGluR5 signaling tunes emotional and cognitive processing in the adult brain

**DOI:** 10.1101/2025.07.07.663457

**Authors:** João Filipe Viana, José Duarte Dias, Candela González-Arias, Luís Samuel Alves, Alexandra Veiga, Daniela Sofia Abreu, João Luís Machado, Sara Barsanti, Rui Jorge Nobre, Luís Pereira de Almeida, Gertrudis Perea, João Filipe Oliveira

## Abstract

The hippocampus is a brain region involved in both emotion regulation and higher cognitive functions. Astrocytes have emerged as active modulators of synaptic activity, capable of sensing, integrating, and responding to neuronal signals. At glutamatergic synapses, astrocytes detect glutamate through the activation of the metabotropic glutamate receptor 5 (mGluR5). However, most existing research has focused on the role of mGluR5 in developing rodents or in pathological contexts, likely because of the reported lower astrocytic mGluR5 expression levels in adulthood compared to postnatal stages. Importantly, prior studies and our preliminary data have demonstrated mGluR5-mediated signaling in astrocytes of adult mice, supporting a role for this receptor. Therefore, the main objectives of this study were (1) to determine whether these lower levels of mGluR5 are sufficient to activate astrocytes in the adult brain and (2) to investigate whether this activation is involved in regulating circuit function and behavior. To address these objectives, we evaluated adult mice employing a combination of calcium imaging in astrocytes, and loss- and gain-of-function manipulations to assess synaptic plasticity and behavior in adult mice.

First, we found that astrocytes of adult mice display fully functional mGluR5-dependent calcium activity. To examine the role of this activity, we induced the deletion of mGluR5 in astrocytes across the entire brain of adult mice. These mice developed anxious- and depression-like behaviors, along with reduced sociability and recognition memory, but showed increased behavioral flexibility. These results highlighted the hippocampus as a key region for mGluR5-mediated astrocytic influence on behavior, leading us to specifically target hippocampal astrocytes. A viral-driven ablation in this area demonstrated that astrocytic mGluR5 plays a role in both basal transmission and the regulation of synaptic plasticity. Behaviorally, the deletion of astrocytic mGluR5 in the hippocampus recapitulated anxious-like behaviors, social deficits, and impaired long-term recognition memory. Surprisingly, it improved place recognition memory but reduced behavioral flexibility. Lastly, overexpressing this receptor to enhance mGluR5 signaling specifically in hippocampal astrocytes impaired place recognition memory but improved behavioral flexibility, revealing a role for astrocytic mGluR5 in regulating these behaviors.

Overall, our results confirmed the biological relevance of astrocytic mGluR5 during adulthood, specifically in modulating hippocampal function.

## Introduction

Throughout the years, evidence has been establishing that astrocytes are key players in maintaining brain homeostasis and modulating synaptic activity, particularly in cortico-limbic circuits, which impacts cognitive processing and emotional regulation (Oliveira et al., 2015; Nagai et al., 2021). In these brain regions, glutamate is the primary excitatory neurotransmitter, as most cells, such as those in the hippocampus, are either glutamatergic neurons or express glutamate receptors (Gasiorowska et al., 2021). To sense glutamatergic activity, astrocytes express glutamate receptors, namely the metabotropic glutamate receptor 5 (mGluR5) (Panatier and Robitaille, 2016; Dias et al., 2025).

The mGluR5 is a G_q_-protein-coupled receptor that induces intracellular Ca^2+^ elevations in astrocytes, which activate Ca^2+^-dependent mechanisms that modulate basal synaptic activity and neuronal synchrony, acting in both pre- and postsynaptic neurons at the hippocampus (Angulo et al., 2004; Fellin et al., 2004; Panatier et al., 2011). Most of the known involvement in synaptic modulation resulting from astrocytic mGluR5 activation stems from studies in the juvenile brain (Araque et al., 2014). We now know that the expression of mGluR5 in astrocytes is developmentally regulated, decreasing throughout the lifetime (Sun et al., 2013). However, it is essential to note that the physiological relevance of mGluR5 may be more closely related to receptor sensitivity and efficiency rather than its expression levels in astrocytes (Panatier and Robitaille, 2016). Indeed, these lower levels of expression are still sufficient (and probably required) to trigger the activation of astrocytes, as shown in previous studies in adult (Wang et al., 2006) and aged mice (Gómez-Gonzalo et al., 2017). Despite the importance of this receptor for circuit function, most available studies on the roles of astrocytic mGluR5 were primarily performed *ex vivo* in young rodents or cell cultures, missing the physiological properties and functional consequences during adulthood, which are crucial for fully understanding glutamatergic signaling in brain circuits in healthy and pathological conditions (Dias et al., 2025).

The main objectives of this study were (1) to determine whether these lower levels of mGluR5 are sufficient to activate astrocytes in the adult brain and (2) to investigate whether this activation is involved in regulating circuit function and behavior. To address these objectives, we use a combination of loss- and gain-of-function manipulations of levels of mGluR5 expression in astrocytes with Ca^2+^-imaging, synaptic plasticity studies and behavioral testing in adult mice.

Our results revealed that astrocytes of adult mice display fully functional mGluR5-dependent calcium activity, and that the manipulation of this receptor modulates cognitive functions and emotional regulation.

## Results

### Astrocytes display mGluR5-dependent calcium activity in adulthood

Firstly, we aimed to confirm that astrocytes present mGluR5-dependent Ca^2+^ events, since some studies report that astrocytes from adult mice express minimal astrocytic mGluR5 and undetectable mGluR5-evoked Ca^2+^ activity (Sun et al., 2013; Umpierre et al., 2019). For this, *ex vivo* Ca^2+^ imaging was performed in acute brain slices from wild-type mice, using the fluorescent Ca^2+^ indicator dye Fluor-4-AM (n = 2, 5-month-old mice) and the genetically encoded Ca^2+^ indicator GCaMP6f (n = 2, 3-month-old mice) (Figure 1A). Here, we demonstrated that naïve astrocytes exhibit Ca^2+^ events in response to DHPG, a mGluR group I agonist (Figure 1B). Indeed, our results show that after DHPG (1 mM) was locally applied by a micropipette, astrocytes exhibited Ca^2+^ events with bigger amplitudes and increased frequency both by using Fluo-4-AM (Figure 1C, Amplitude: *p* = 0.007, Frequency: *p* = 0.0006) and GCaM6f (Figure 1D, Amplitude: *p* = 0.04, Frequency: *p* = 0.02). These readouts confirm the existence of functional astrocytic mGluR5, indicating that mature astrocytes also display mGluR5-evoked Ca^2+^ elevations. Moreover, since all recordings were performed in the presence of mGluR1 antagonist, it ensures that DHPG is only activating mGluR5.

**Figure 1.**
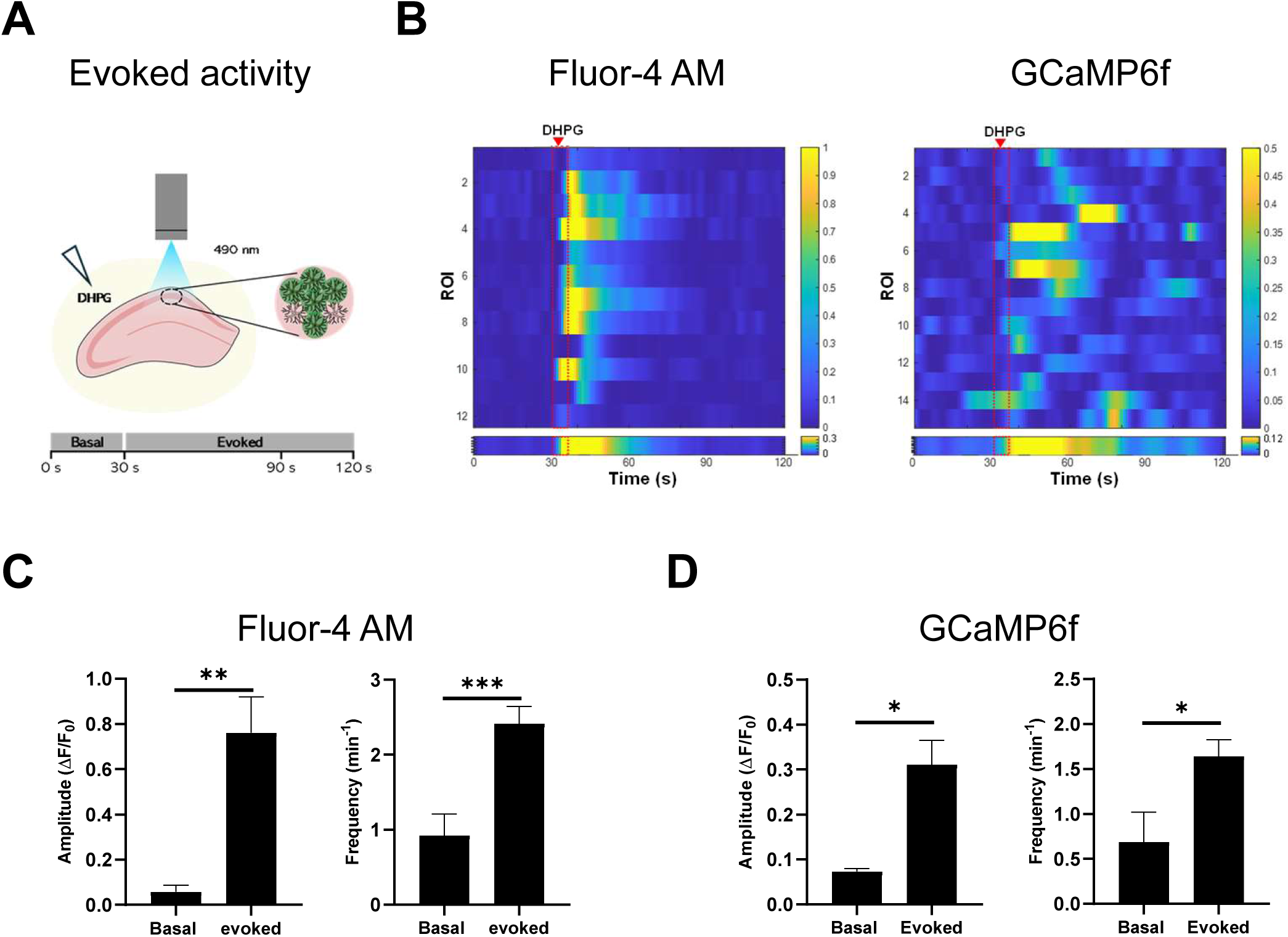
Astrocytes from adult wild-type mice still exhibit mGluR5-dependent calcium activity. (A-D) *Ex vivo* calcium (Ca^2+^) imaging in the CA1 region of acute hippocampal slices from wild-type mice using Fluor-4-AM and GCaMp6f Ca^2+^ indicators. (A) Schematic representation of DHPG-evoked Ca^2+^ events in hippocampal astrocytes. (B) Heatmaps of DHPG-evoked ROIs activity (top) and average population activity (bottom) of Fluor-4-AM (left) and GCaMP6f (right). Color code denotes fluorescence changes. (C-D) Graphical representations of Ca^2+^ events amplitude and frequency of astrocytes using Fluor-4-AM (n = 2 mice, 12 ROIs) and GCaMP6f (n = 2 mice, 15 ROIs). Data plotted as mean ± SEM and analyzed using Student’s t-test, being * *p* < 0.05, ** *p* < 0.01 and *** *p* < 0.001.

### ALDH1L1-mGluR5KO mice exhibit decreased levels of mGluR5 in hippocampal astrocytes

A tamoxifen-inducible recombination strategy was employed to induce the deletion of the mGluR5 gene in ALDH1L1-positive cells. To evaluate the efficiency of the tamoxifen protocol, four weeks after the last injection, ALDH1L1-mGluR5KO mice (n = 3) were euthanized, followed by brain removal and sectioning to assess tdTomato reporter protein expression throughout the brain, which indicates genetic recombination. We observed tdTomato expression throughout the brain (Figure 2A), indicating efficient genetic recombination. Furthermore, these tdTomato-positive cells exhibited the typical star-shaped morphology of astrocytes, with no evidence of other cell-type morphologies. To confirm that recombination occurred specifically in astrocytes, an immunohistochemistry protocol was used to label different cell types in the hippocampus of ALDH1L1-mGluR5KO mice and evaluate co-expression with tdTomato-expressing cells. Then, the specific marks GFAP and S100β were used to label astrocytes, while NeuN, Iba1, and CC-1 were used to label neurons, microglia, and oligodendrocytes, respectively (Figure 2B-F). Our results show that tdTomato co-localizes with cells expressing both GFAP (Figure 2B) and S100β (Figure 2C), which confirms the recombination in astrocytes. In addition, no co-expression with NeuN (Figure 2D), Iba1 (Figure 2E), and CC-1 (Figure 2F) was observed in cells expressing these proteins, excluding any recombination in neurons, microglia, and oligodendrocytes, respectively. Thus, the results confirm that genetic recombination occurred specifically in astrocytes within the hippocampus. To confirm the ablation of mGluR5 in astrocytes upon tamoxifen-induced genetic recombination, mGluR5 expression levels were quantified by immunofluorescence in the territory of astrocytes in brain slices from Control (n = 3 mice, 9 astrocytes, Figure 2G) and ALDH1L1-mGluR5KO (n = 3 mice, 9 astrocytes, Figure 2H) mice. We observed that ALDH1L1-mGluR5KO mice had lower average intensity values in astrocytes compared to those from Control mice (Figure 2I, p = 0.006), confirming the deletion of mGluR5.

**Figure 2.**
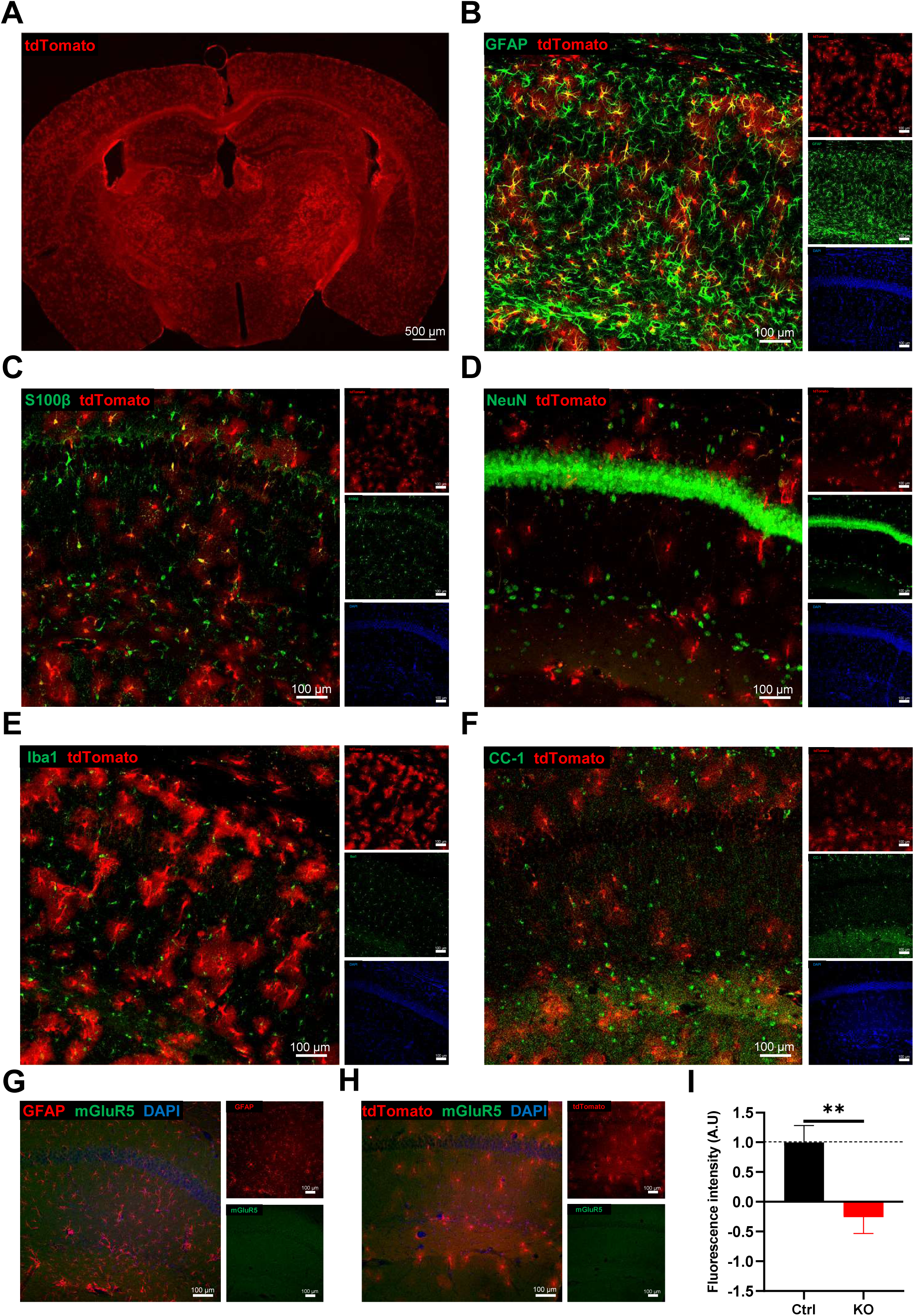
tdTomato expression was specifically for astrocytes in ALDH1L1-mGluR5KO mice and exhibited reduced immunolabeled mGluR5 fluorescence intensity in their territory in the CA1 region of the hippocampus. (A) Representative image (4x) of a hippocampal slice of ALDH1L1-mGluR5KO mouse showing tdTomato expression. (B-F) Representative image (20x) of the CA1 region of the dorsal hippocampus of ALDH1L1-mGluR5KO mouse following immunolabeling of tdTomato (red), DAPI (blue), GFAP (green in B), S100β (green in C), NeuN (green in D), Iba1 (green in E), and CC-1 (green in F). In yellow, it represents co-labeling of different markers in the same cell. (G-H) Representative images (20x) of the CA1 region of the dorsal hippocampus of (G) Control and (H) ALDH1L1-mGluR5KO (C) mice following immune labeling of mGluR5 (green), DAPI (blue) and tdTomato (red). Scale bars are depicted in the image. (I) Mean intensity of fluorescence quantification of immunolabeled mGluR5 within the astrocytic GFAP territory. Data was plotted as mean ± SEM and analyzed using an independent t-test, being ** *p* < 0.01. Astrocytes from Control (Ctrl) are plotted in a black bar (n = 9), and astrocytes from ALDH1L1-mGluR5KO (KO) mice are plotted in a red bar (n = 9).

Overall, the results demonstrate that genetic recombination specifically occurred in astrocytes and induced a reduction in astrocytic mGluR5 expression, validating the use of this mouse model to study the importance of astrocytic mGluR5 in brain processing.

### Deletion of astrocytic mGluR5 promotes anxious- and depressive-like behaviors in adulthood

To understand if mGluR5 is involved in anxious-like behavior, the ALDH1L1-mGluR5KO (n_males_ = 25 and n_females_ = 17) and respective control (n_males_ = 18 and n_females_ = 20) mice were tested in the L/D box, a test based on the natural preference of mice to dark and enclosed spaces (Figure 3A). Our results showed that ALDH1L1-mGluR5KO mice from both sexes spent less time in the light zone when compared to the respective control mice (Figure 3A, *p* = 0.02). In addition, male and female Control mice behaved similarly as expected (Figure 3A). Furthermore, to continue exploring the role of astrocytic mGluR5 in mood regulation, ALDH1L1-mGluR5KO (n_males_ = 30 and n_females_ = 20) and respective control (n_males_ = 28 and n_females_ = 19) were subjected to the TST to assess learned helplessness, a type of depressive-like behavior, measured by states of immobility in unescapable situations (Figure 3B). In the TST, we observed that male and female ALDH1L1-mGluR5 mice exhibited higher immobility compared to their respective control mice (Figure 3B, *p*_male_ = 0.006, *p*_females_ = 0.01). Finally, ALDH1L1-mGluR5KO mGluR5KO (n_males_ = 30 and n_females_ = 16) and respective control (n_males_ = 27 and n_females_ = 18) performed the FST (Figure 3C). Similarly to the TST results, ALDH1L1-mGluR5KO mice from both sexes exhibited higher immobility times than their respective Control mice (Figure 3C, *p*_male_ = 0.02, *p*_females_ = 0.05). Moreover, no differences were found when comparing male and female Control mice (Figure 3B-C). After assessing the anxious- and depressive-like behavior of ALDH1L1-mGluR5KO mice following astrocytic mGluR5 deletion, we proceeded to study its implications in social behavior, a behavioral dimension often disrupted by mood disorders. Hence, ALDH1L1-mGluR5KO (n_males_ = 9-10 and n_females_ = 9-10) and respective control mice (n_males_ = 9-10 and n_females_ = 8-10) performed the 3CST. Firstly, mice performed the 3CST-SP test (Figure 3D), in which we observed that male ALDH1L1-mGluR5KO mice spent less time interacting with the stranger mouse (Figure 3D, *p* = 0.04), Interestingly, both male and female ALDH1L1-mGluR5KO mice exhibited a lower S.I. (Figure 3D, *p*_male_ = 0.03, *p*_female_ = 0.004) compared to the respective controls. Moreover, in the 3CST-SM test (Figure 3E), ALDH1L1-mGluR5KO mice exhibited intact social memory compared to control mice, as indicated by the similar amount of time spent interacting with the novel mouse (Figure 3E) and SI (Figure 3E).

**Figure 3.**
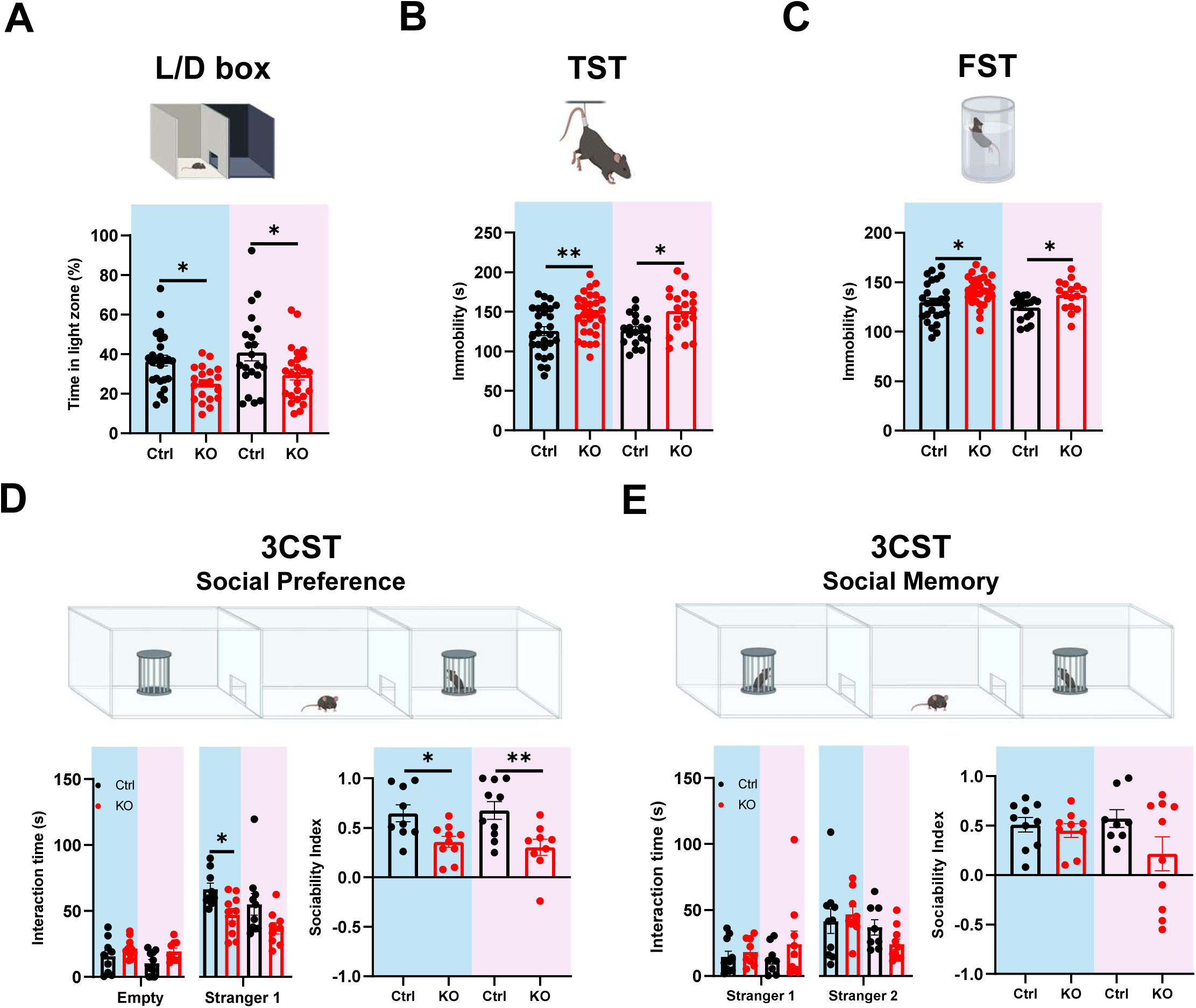
Deletion of astrocytic mGluR5 induces anxious-like behavior, learned helplessness, and reduced sociability in ALDH1L1-mGluR5KO mice. (A) Light/Dark (L/D) box test illustration and graphical representation of the percentage of time spent in the light zone of the L/D Box apparatus. (B) Tail Suspension Test (TST) illustration and graphical representation of time immobile in the TST. (C) Forced Swin Test (FST) illustration and graphical representation of immobility time in the FST. (D) Three Chambers Social Test - Social Preference (3CST-SP) illustration and Graphical representation of the time interacting with the empty cup and Stranger 1 (left) and respective Sociability Index (S.I.) (right). (E) Three Chambers Social Test - Social Memory (3CST-SM) illustration and graphical representation of the time interacting with Stranger 1 and Stranger 2 (left) and respective Sociability Index (S.I.) (right). Data plotted as mean ± SEM and analyzed using (D-E) Student’s t-test or Three-way ANOVA (A-E), being * *p* < 0.05 and ** *p* < 0.01. Male mice are highlighted in blue and female mice in pink. Control (Ctrl) mice are plotted in a black bar, and individual values in black dots (n = 8-28), and ALDH1L1-mGluR5KO (KO) mice are plotted in a red bar, and individual values in red dots (n = 9-30).

Overall, our results suggest that astrocytic mGluR5 is important for emotional regulation, specifically being involved in anxious- and depressive-like behaviors, also compromising sociability in mice. These results persuaded us to investigate the involvement of the astrocytic receptor in cognitive function.

### Deletion of astrocytic mGluR5 leads to deficits in long-term memory and enhances behavioral flexibility

To determine if astrocytic mGluR5 is involved in cognitive processing, ALDH1L1-mGluR5KO mice and their respective controls were tested in various behavioral paradigms, including the 2TPR to assess place recognition, NOR to assess object recognition memory, and the MWM to assess spatial reference memory and behavioral flexibility.

First, ALDH1L1-mGluR5KO (n_males_ = 22 and n_females_ = 12) and Control (n_males_ = 16 and n_females_ = 9) mice were tested in the 2TPR test. The test was used to assess place recognition memory, taking advantage of mice’s natural drive to explore novelty (Figure 4A). The obtained results indicate that ALDH1L1-mGluR5KO mice, similarly to the controls, were able to discriminate the novel arm from the previously explored arms, as shown by the D.I. related to time spent exploring the novel arm (Figure 4A). Following the 2TPR, ALDH1L1-mGluR5KO mice (n_males_ = 11 and n_females_ = 11-12) and their respective controls (n_males_ = 10-13 and n_females_ = 10) mice were tested using the NOR task. To assess recognition memory, mice underwent either a 1 h retention interval for short-term memory (Figure 4B) or a 24 h interval for long-term memory (Figure 4C) before exposure to the novel object. In the 1 h retention interval task, mice from all experimental groups presented similar D.I. related to the time interacting with the novel object vs. the familiar object (Figure 4B). However, when the retention interval was increased to 24 h, male and female ALDH1L1-mGluR5KO mice showed lower D.I. compared to control mice (Figure 4C, *p*_male_ = 0.003, *p*_females_ = 0.002). Additionally, no differences were found between male and female mice (Figure 4A-C). To further explore the role of astrocytic mGluR5 in cognition, ALDH1L1-mGluR5KO (n_males_ = 24-26 and n_females_ = 14-22) and control (n_males_ = 28 and n_females_ = 12-18) mice were tested in the MWM. This test was divided into two tasks, namely, the spatial reference task (Figure 4D) and the reversal learning task (Figure 4E). In the spatial reference task, mice had to learn the location of a hidden platform, kept in a fixed position across all days, by relying on spatial cues. Our results show that mice from all experimental groups successfully learned to find the platform, showing similar escape latency and distance swam (Figure 4D, *p* < 0.0001). On the fifth day, the mice performed the reversal learning task, in which the platform was relocated to a new quadrant while the spatial cues remained unchanged. Mice from all experimental groups learned the new platform location, spending more time and swimming greater distances in the new quadrant compared to the previous platform quadrant (Figure 4E, *p* < 0.01). Intriguingly, male ALDH1L1-mGluR5KO mice spent even more time in the new quadrant than Control mice (Figure 4E, *p* = 0.02), showing enhanced behavioral flexibility.

**Figure 4.**
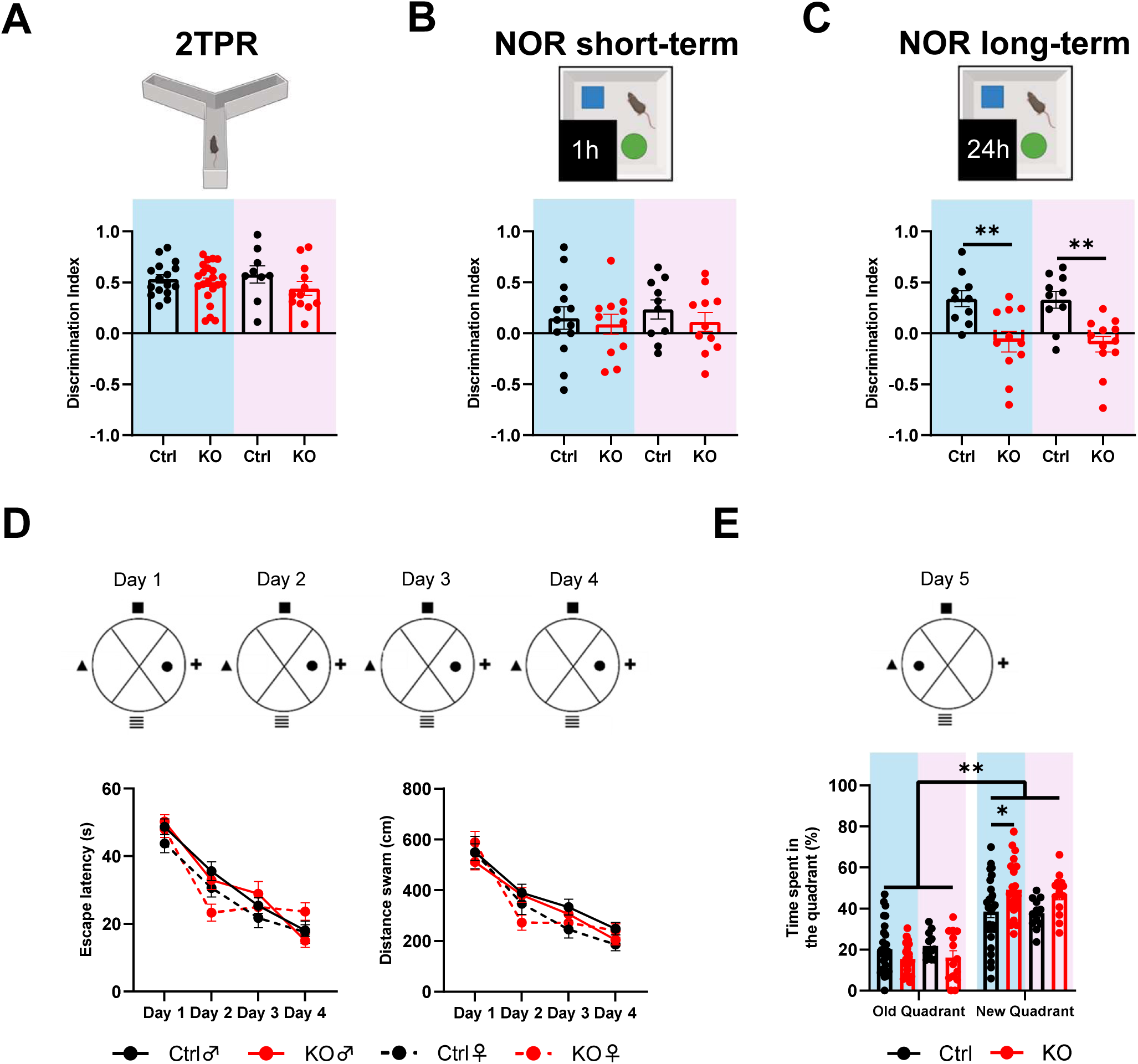
Ablation of mGluR5 in astrocytes impairs long-term recognition memory but enhances behavioral flexibility. (A) Two-trial Place Recognition (2TPR) illustration and graphical representation of the discrimination Index (D.I.) for time spent exploring the novel arm vs. familiar and start arms. (B) Novel Object Recognition (NOR) to assess short-term memory illustration and graphical representation of the discrimination Index (D.I.) for time interacting with the new vs. familiar objects. (C) NOR to test long-term memory illustration and graphical representation of the discrimination Index (D.I.) for time interacting with new vs. familiar objects. (D) Spatial reference memory task of the Morris Water Maze (MWM) illustration and respective Learning Curves for escape latency (left) and distance swam (right). (E) Graphical representation of the time spent swimming in the old and new quadrants during the Reversal learning task. Data plotted as mean ± SEM and analyzed using Student’s t-test, being * *p* < 0.05, ** *p* < 0.01. Male mice are highlighted in blue and female mice in pink. Control (Ctrl) mice are plotted in a black bar and individual values in black dots (n = 9-28) and ALDH1L1-mGluR5KO (KO) mice are plotted in a red bar and individual values in red dots (n = 11-26).

Overall, the results indicate that astrocytic mGluR5 is involved in cognitive processing, specifically in long-term recognition memory and behavioral flexibility. Our findings prompted us to continue exploring the biological relevance of astrocytic mGluR5 behavior, with a specific focus on the hippocampus. We focus on the hippocampus due to its crucial role in emotional regulation and cognition. Additionally, most of the behavioral tasks used were selected based on their dependence on the hippocampus.

### Hippocampal manipulation of astrocytic mGluR5 expression tunes calcium responsiveness in adulthood

To induce mGluR5 knockout specifically in astrocytes of the dorsal hippocampus, a viral approach was employed following protocols previously validated in our laboratory to grant viral transduction and cellular specificity. Thus, to validate mGluR5 deletion, specifically in astrocytes, we performed a combination of IHC-based fluorescence quantification for protein expression levels and *ex vivo* Ca^2+^ imaging for functional confirmation (Figure 5). First, astrocytic mGluR5 expression levels were quantified by immunolabeling mGluR5 in brain tissue of dHIP:GFAP-mGluR5KO (n = 3) and respective Control (n = 3) mice. Moreover, the GFAP marker was used to identify astrocytes while the mCherry reporter gene was used to label cells that underwent genetic recombination (Figure 5A-C). Our quantification results showed that brain slices from dHIP:GFAP-mGluR5KO mice exhibited decreased average intensity values within the territory of mCherry-positive astrocytes when compared to astrocytes in brain slices of Control mice (Figure 5C, *p* < 0.0001). Then, to confirm functional mGluR5 deletion in astrocytes *ex vivo*, Ca^2+^ imaging was performed in hippocampal brain slices of dHIP:GFAP-mGluR5KO (n = 3) and respective Control (n = 2) mice. Therefore, brain slices were incubated with the Ca^2+^ indicator dye Fluor-4-AM, and Ca^2+^ events were recorded in response to the local application of DHPG, an agonist of the mGluR5 receptor (Figure 5D-E). Our results show that in control animals, when DHPG (1 mM) was locally applied by a micropipette, astrocytes displayed Ca^2+^ events with larger amplitudes (Figure 5F, *p* = 0.03) and higher frequencies (Figure 5F, *p* = 0.0003) Ca^2+^ events when compared to basal conditions. Secondly, the same analysis was performed in brain slices of dHIP:GFAP-mGluR5KO. When stimulated by DHPG application, we observed that mGluR5KO astrocytes (n = 30) presented reduced Ca^2+^ events (Figure 5E), as shown by no alteration in amplitude and frequency (Figure 5G). This indicates that the deletion of mGluR5 was efficient in downregulating Ca^2+^ signaling. Altogether, our results confirm that we successfully knocked out mGluR5 specifically in astrocytes, validating the use of the dHIP:GFAP-mGluR5KO mouse model to study mGluR5 in the hippocampus.

**Figure 5.**
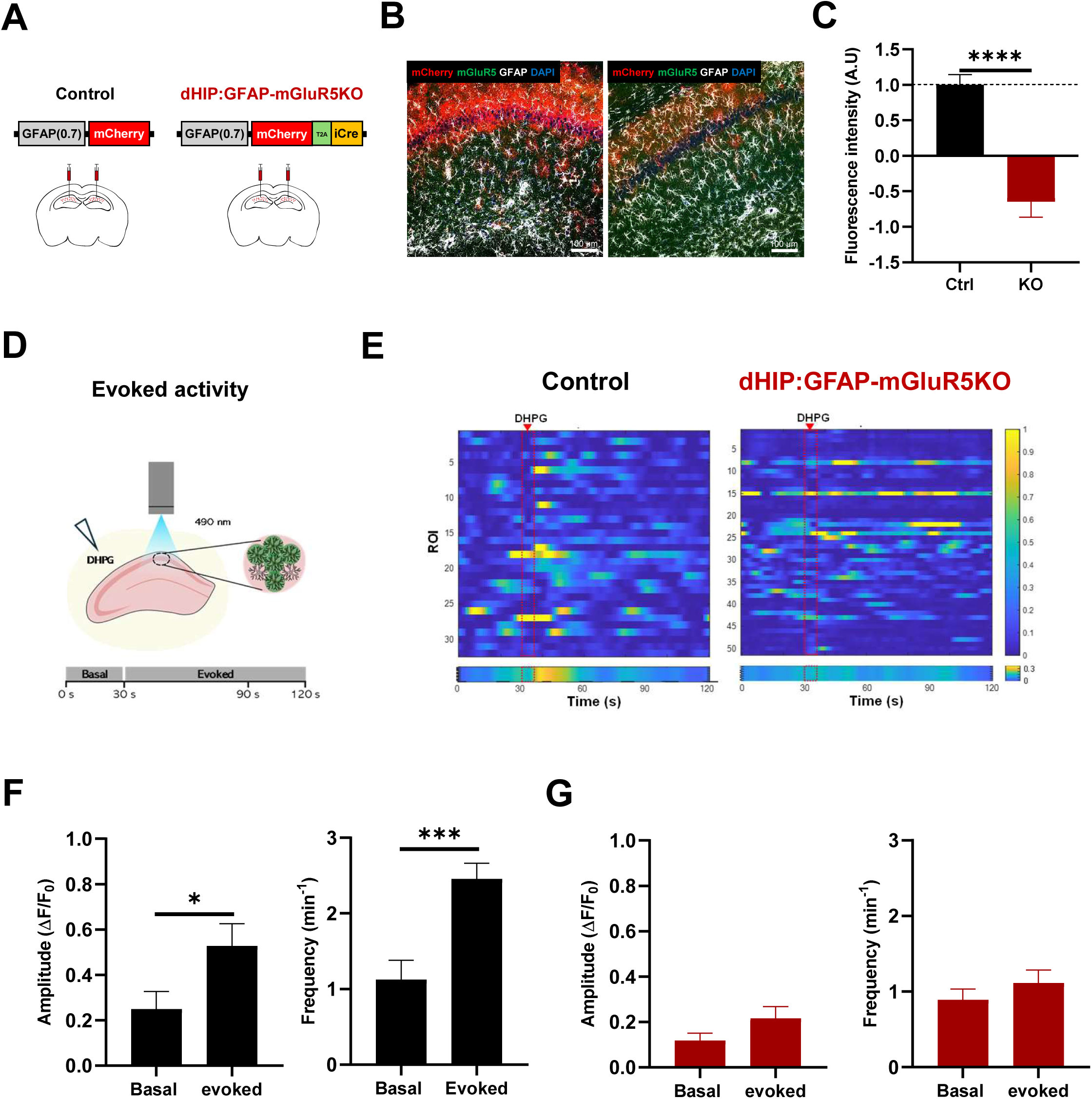
AAV-transduced astrocytes present reduced mGluR5 fluorescence intensity and abolished DHPG-evoked Ca^2+^ events in CA1 *stratum oriens* and *stratum radiatum* of the dorsal hippocampus. (A) Representative scheme depicting the bilateral stereotaxic injections of AAV5:GFAP-mCherry and AAV5:GFAP(0.7)-mCherry-T2A-iCre virus in the hippocampal CA1 area of mGluR5^flox/flox^ mice. (B) Representative images of the CA1 region of the dorsal hippocampus of Control (left) and dHIP:GFAP-mGluR5KO (right) mice after immunolabeling of mGluR5 (green), GFAP (white), mCherry (red), DAPI (blue). Scale bar is depicted in the image. (C) mGluR5 mean intensity of fluorescence quantification within the astrocytic GFAP-defined territory (n_Control_ = 22, n_dHIP:GFAP-mGluR5KO_ = 24). (D-G) Ex vivo calcium (Ca^2+^) imaging in the CA1 region of acute hippocampal slices from Control and dHIP:GFAP-mGluR5KO mice. (D) Schematic representation of DHPG-evoked Ca2+ events in hippocampal astrocytes. (E) Heatmaps of DHPG-evoked ROIs activity (top) and average population activity (bottom) of Control (left) and dHIP:GFAP-mGluR5KO (right) mice. Color code denotes fluorescence changes. (F-G) Graphical representations of Ca^2+^ events amplitude (left) and frequency (right) of astrocytes from Control (F, n = 2 mice, 32 ROIs) and dHIP:GFAP-mGluR5KO (G, n = 3 mice, 65 ROIs) mice. Data plotted as mean ± SEM and analyzed using Student’s t-test, being * *p* < 0.05, ** *p* < 0.01, and **** *p* < 0.0001. Control (Ctrl) mice are plotted in a black bar, and dHIP:GFAP-mGluR5KO (KO) mice are plotted in a red bar.

Alternatively, we overexpress mGluR5 in astrocytes in the adult hippocampus, by using a viral vector, AAV5:GFAP(0.7)-FLAG-shortGrm5, injected in the dorsal hippocampus of wildtype mice. Four weeks later, the receptor overexpression was confirmed by fluorescence quantification of mGluR5 expression and validated by *ex vivo* Ca^2+^ imaging (Figure 6). The viral vector used to overexpress mGluR5 also induced the expression of a FLAG tag, allowing for the identification of transduced cells. However, due to technical constraints, it was not possible to immunolabel the FLAG tag. This was likely due to the expression of a single FLAG tag, which has been described as potentially lacking sufficient epitope density for IHC detection (Ferrando et al., 2015). Despite these limitations, the mGluR5 fluorescence quantification was performed by IHC to label mGluR5 and GFAP in dHIP:GFAP-mGluR5+ (n = 3) and Control (n = 3) mice, revealing an increase in mGluR5 fluorescence compared with controls, in which astrocytic mGluR5 levels are reported to be low, but still a detectable fluorescence signal (Figure 6A-C). In fact, when mGluR5 fluorescence was quantified in the territory of GFAP-positive cells from brain slices of dHIP:GFAP-mGluR5+ (n = 3) and Control (n = 3) mice, we observed higher average mGluR5 fluorescence intensity in astrocytes of the dorsal hippocampus of dHIP:GFAP-mGluR5+ mice, confirming mGluR5 overexpression (Figure 6C, *p* < 0.0001). Then, to determine if the overexpression of mGluR5 modified astrocytic Ca^2+^ activity, we performed functional studies by *ex vivo* Ca^2+^ imaging, taking advantage of the genetically encoded Ca^2+^ indicator GCaMP6f. Astrocyte Ca^2+^ events were monitored in the CA1 *stratum oriens* and *stratum radiatum* of the dorsal hippocampus of dHIP:GFAP-mGluR5+ (n = 3) and respective Control (n = 3) mice. As in the Fluo-4 experiments, DHPG was locally applied through a micropipette to stimulate Ca^2+^ signaling in astrocytes from dHIP:GFAP-mGlu5+ (n = 154 ROIs) and Control (n = 73 ROIs) mice (Figure 6D-G). Our results showed that DHPG evoked Ca^2+^ events with bigger amplitude (Figure 6F, Control: *p* = 0.004 and Figure 6G, mGluR5+: *p* < 0.0001) and frequency (Figure 6F, Control: *p* < 0.0001 and Figure 6G, mGluR5+: *p* < 0.0001) in astrocytes from mice of both experimental groups. Remarkably, DHPG-evoked Ca^2+^ events in astrocytes from dHIP:GFAP-mGluR5+ mice presented increased amplitude compared to basal activity, but also when comparing DHPG-mediated Ca^2+^ dynamics in astrocytes from Control mice. Interestingly, the frequency appears to be lower in astrocytes from dHIP:GFAP-mGluR5+. Overall, these results confirm the functional overexpression of mGluR5 in astrocytes, validating the dHIP:GFAP-mGluR5+ mouse model to study the enhancement of mGluR5 signaling in astrocytes.

**Figure 6.**
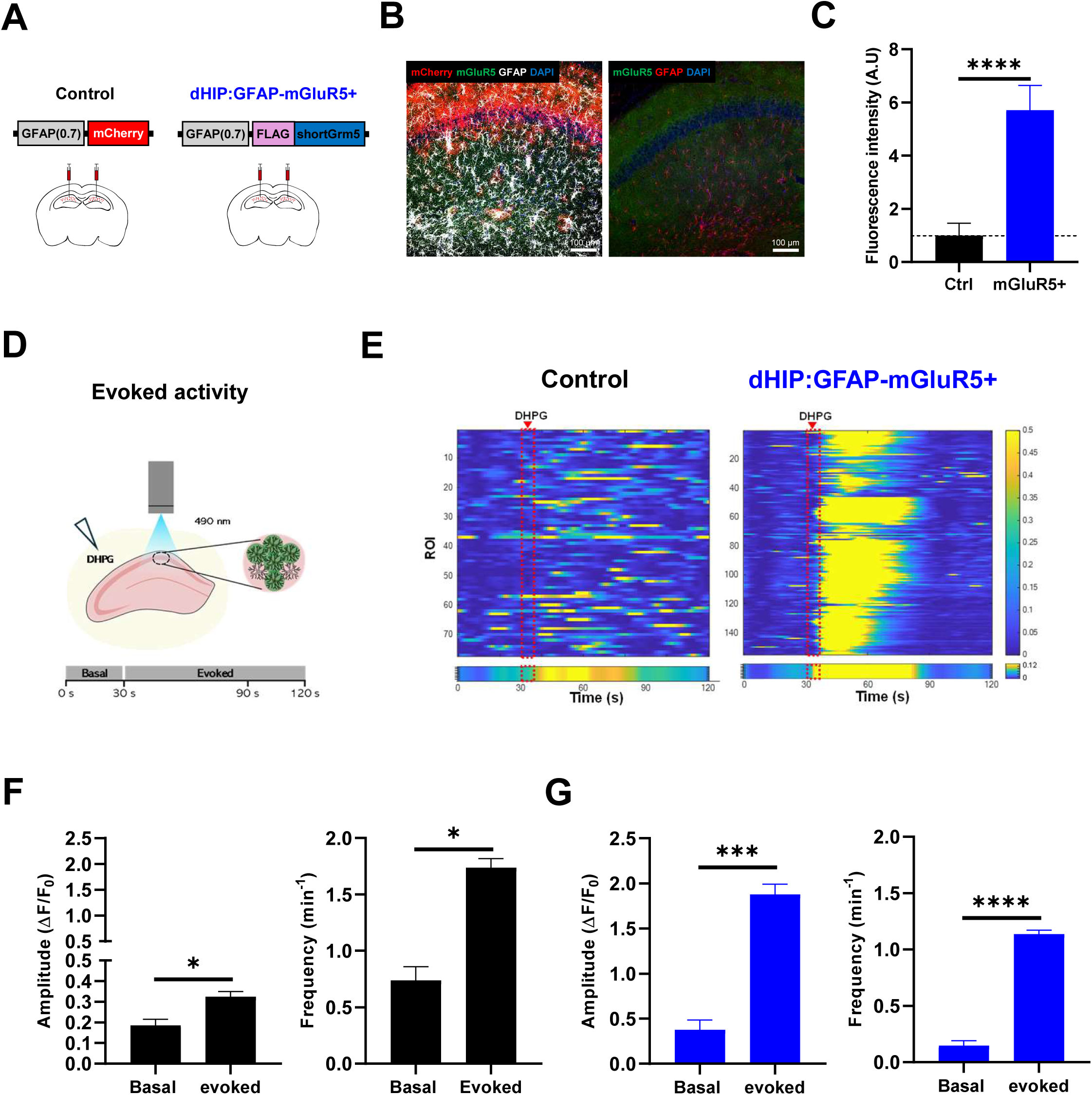
Overexpression of mGluR5 in astrocytes in CA1 *stratum oriens* and *stratum radiatum* of the dorsal hippocampus increases mGluR5 fluorescence intensity and enhances DHPG-evoked Ca^2+^ events in astrocytes. (A) Representative scheme depicting the bilateral stereotaxic injections of AAV5:GFAP-mCherry and AAV5:GFAP(0.7)-FLAG-shortGrm5 virus in the hippocampal CA1 area of C57BL/6 mice. (B) Representative images of the CA1 region of the dorsal hippocampus of Control (left) and dHIP:GFAP-mGluR5+ (right) mice after immunolabeling of mGluR5 (green), GFAP (white), mCherry (red), DAPI (blue). Scale bar is depicted in the image. (C) mGluR5 mean intensity of fluorescence quantification within the astrocytic GFAP-defined territory (n_Control_ = 21, n_dHIP:GFAP-mGluR5KO_ = 22). (D-G) Ex vivo calcium (Ca^2+^) imaging in the CA1 region of acute hippocampal slices from Control and dHIP:GFAP-mGluR5+ mice. (D) Schematic representation of DHPG-evoked Ca2+ events in hippocampal astrocytes. (E) Heatmaps of DHPG-evoked ROIs activity (top) and average population activity (bottom) of Control (left) and dHIP:GFAP-mGluR5+ (right) mice. Color code denotes fluorescence changes. (G-F) Graphical representations of Ca^2+^ events amplitude (left) and frequency (right) of astrocytes from Control (F, n = 3 mice, 75 ROIs) and dHIP:GFAP-mGluR5+ (G, n = 3 mice, 154 ROIs) mice. Data plotted as mean ± SEM and analyzed using Student’s t-test, being ** *p* < 0.01, *** *p* < 0.001, and **** *p* < 0.0001. Control (Ctrl) mice are plotted in a black bar, and dHIP:GFAP-mGluR5+ (mGluR5+) mice are plotted in a blue bar.

### Bidirectional regulation of synaptic plasticity by astrocytic mGluR5 in the dorsal hippocampus

To further understand the role of astrocytic mGluR5 in the modulation of synaptic transmission, we first analyzed the I/O responses in acute hippocampal slices from Control (n = 3 mice, 9 slices) and dHIP:GFAP-mGluR5KO (n = 4 mice, 8 slices) mice (Figure 7A-B). Our results indicate that the deletion of astrocytic mGluR5 increased the synaptic response to higher-intensity stimulations, specifically at 1000 mA (*p* = 0.001) and 1300 mA (*p* = 0.0009), compared to the Control (Figure 7B), indicating enhanced excitability. Then, to pursue investigating how astrocytic mGluR5 affects synaptic plasticity in the hippocampus, a theta-burst protocol was applied to the synaptic inputs, CA3-CA1 synaptic pathway, to induce LTP in acute hippocampal slices from Control (n = 3 mice, 5 slices) and dHIP:GFAP-mGluR5KO (n = 4 mice, 6 slices) mice. The theta-burst stimulation led to an initial enhancement of the fEPSP slope in Control mice, which increased over time, followed by a stabilization period until the end of the recording (Figure 7C-D). However, hippocampal slices of dHIP:GFAP-mGluR5KO mice stimulated with this protocol showed a slight increase in fEPSP slope, which began to decrease shortly and returned to baseline values by the end of the recording(Figure 7C-D). Indeed, after fifty minutes of recording, LTP magnitude was smaller in dHIP:GFAP-mGluR5KO mice when compared to Control mice (Figure 7E, *p* = 0.01), suggesting the role of astrocytic mGluR5 as a modulator of adult hippocampal LTP. After understanding how astrocytic mGluR5 is involved in basal synaptic transmission and LTP, we performed the same experiment in acute hippocampal slices from dHIP:GFAP-mGluR5+ mice (n = 5 mice, 6-12 slices) to investigate how increased expression of m-GluR5 influences these phenomena. Firstly, through the I/O curve assessment, we verified that overexpression of mGluR5 in astrocytes did not alter basal synaptic transmission compared to Control mice (Figure 7B). Furthermore, when synaptic plasticity in the form of LTP was evaluated by theta-burst stimulation, mice overexpressing mGluR5 displayed a decrease of fEPSP slope that was maintained over time until the end of the recording (Figure 7C-D). Indeed, after fifty minutes of recording dHIP:GFAP-mGluR5+ slices presented smaller fEPSP slope magnitude when compared to Control mice (Figure 7E, *p* < 0.0001), which denotes excitatory synaptic depression. Altogether, our findings showed that in the adult hippocampus, astrocytic mGluR5 is indeed important for basal synaptic activity and LTP synaptic plasticity, whereas in the case of astrocytic mGluR5 overexpression, it led to long-lasting synaptic depression.

**Figure 7.**
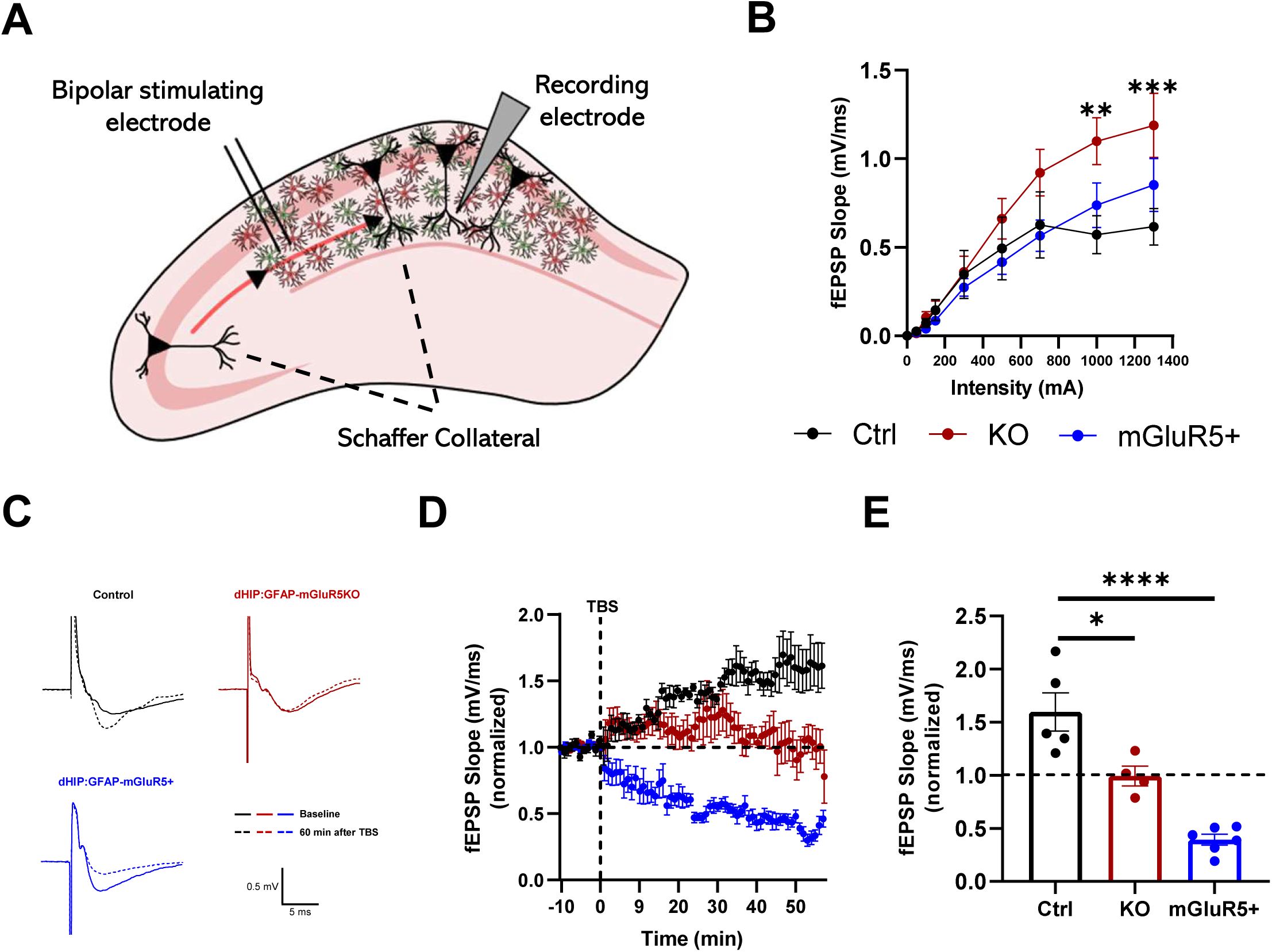
Deletion of astrocytic mGluR5 in the dorsal hippocampus enhances basal synaptic transmission and abolishes long-term potentiation, while overexpression of mGluR5 in astrocytes induces synaptic activity depression. (A) Representative illustration of local field recording of fast excitatory postsynaptic potentials (fEPSPs) of pyramidal neurons in CA1 stratum radiatum of acute hippocampal slices (B) I/O curves depicting fEPSP slope changes with different stimulation intensities in hippocampal slices of Control (n = 3 mice, 9 slices), dHIP:GFAP-mGluR5KO (n = 4 mice, 8 slices), and dHIP:GFAP-mGluR5+ (n = 5 mice, 12 slices). (C) Representative fEPSC traces (average from 20 consecutive responses) recorded before (left) theta-burst stimulation and after 1 hour (right) for Control (top), dHIP:GFAP-mGluR5KO (middle), and dHIP:GFAP-mGluR5+ (bottom). (D) Average of normalized fEPSP slope throughout time before and 60 min after theta-burst stimulation and (E) comparison of the magnitude of LTP in the last 10 minutes of recording in acute hippocampal brain slices from Control (n = 3 mice, 5 slices), dHIP:GFAP-mGluR5KO (n = 4 mice, 6 slices) and dHIP:GFAP-mGluR5+ (n = 3 mice, 6 slices). Data plotted as mean ± SEM and analyzed using (B, D) Two-way ANOVA and (E) One-way ANOVA, being * *p* < 0.05 and **** *p* < 0.0001. Control (Ctrl) mice are plotted in a black bar/dots, dHIP:GFAP-mGluR5KO (KO) mice are plotted in a red bar/dots and dHIP:GFAP-mGluR5+ (mGluR5+) mice are plotted in a blue bar/dots.

### Deletion of mGluR5 in hippocampal astrocytes recapitulated anxious-like behaviors, social deficits, and impaired long-term recognition memory

To investigate how astrocytic mGluR5 in the hippocampus is involved in anxious-like behavior, Control (n = 21) and dHIP:GFAP-mGluR5KO (n = 23) mice performed the LD box test (Figure 8A), which is based on mice’s propensity to prefer dark and enclosed spaces. Our results revealed that dHIP:GFAP-mGluR5KO mice spent less time in the light zone when compared to Control mice (Figure 8A, *p* = 0.04), suggesting the development of an anxious-like phenotype. Next, Control (n = 16) and dHIP:GFAP-mGluR5KO (n = 19) mice were tested in the TST to assess learned despair (Figure 8B). The results obtained show that mice from both experimental groups exhibited similar immobility time (Figure 8B), meaning that mGluR5 in the hippocampus is not involved in learned helplessness. After assessing emotional behavior, Control (n = 12) and dHIP:GFAP-mGluR5KO (n = 12) mice were tested in the 3CST to assess sociability, in 3CST-SP (Figure 8C), and social memory, in 3CST-SM (Figure 8D). In the 3CST-SP, our results indicate that dHIP:GFAP-mGluR5KO mice spend more time interacting with the empty cup than Control mice (Figure 8C, *p* = 0.008), which resulted in a smaller S.I. in dHIP:GFAP-mGluR5KO mice when compared to Control mice (Figure 8C, *p* = 0.01), suggesting social deficits upon mGluR5 deletion. Finally, in the 3CST-SM mice from Control and dHIP:GFAP-mGluR5KO experimental groups, spend a similar amount of time interacting with Stranger 1 (familiar mouse) and then with Stranger 2 (novel mouse) (Figure 8D), which resulted in a similar and positive D.I. between mice from both groups (Figure 8D).

**Figure 8.**
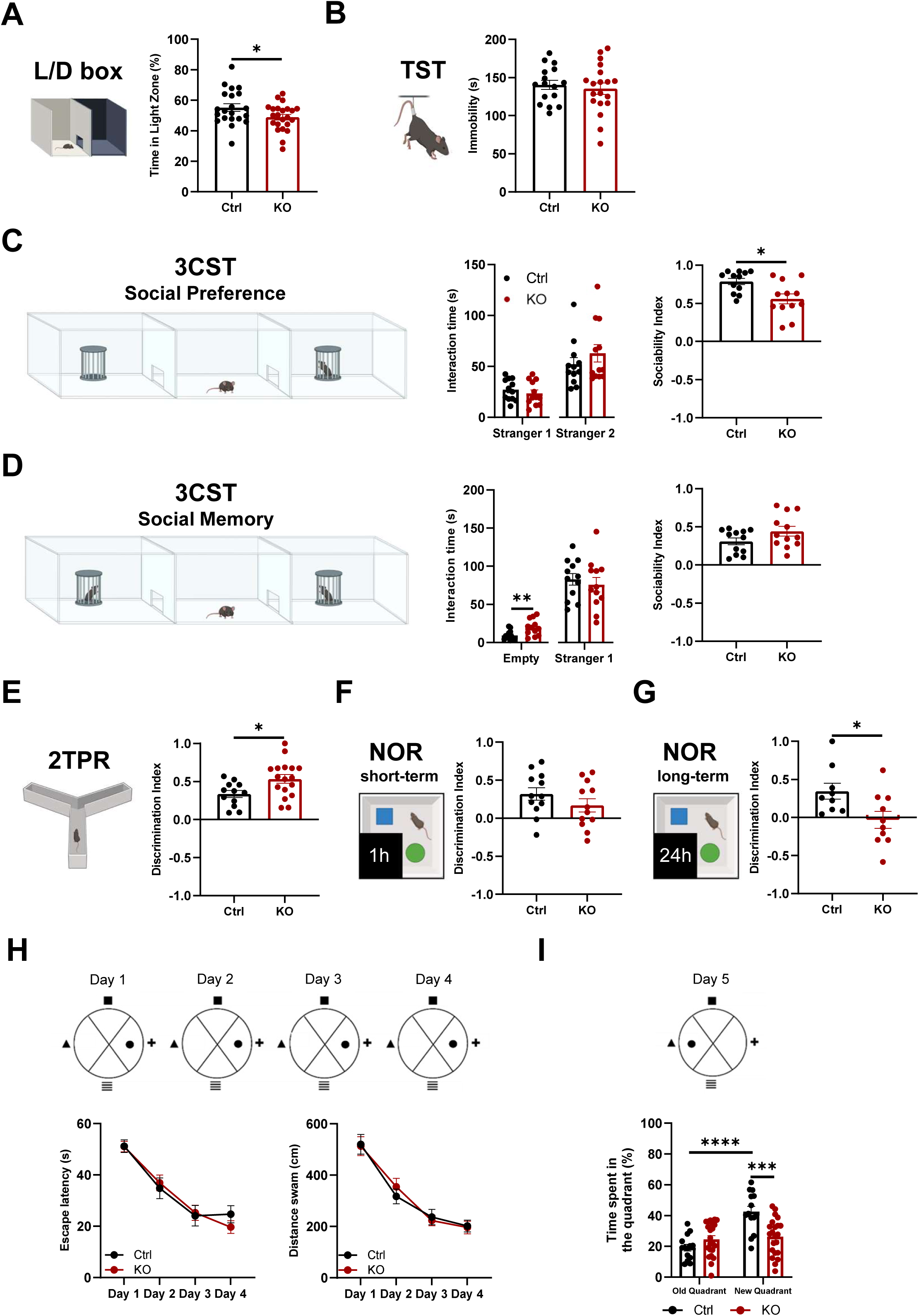
Deletion of mGluR5 in hippocampal astrocytes induces anxious-like behavior, social deficits, and impairs long-term recognition memory and behavioral flexibility but enhances place recognition memory. (A) Representative illustration of the Light/Dark (L/D) box test and percentage of time spent in the light zone of the L/D Box apparatus. (B) Representative Illustration of the Tail Suspension Test (TST) and immobility time in the TST. (C) Representative illustration of the 3 Chambers Social Test - Social Preference (3CST-SP), time interacting with the empty cup and Stranger 1 (left), and Sociability Index (S.I.) for preference to interact with Stranger 1 (right). (D) Representative illustration of the 3 Chambers Social Test - Social Memory (3CST-SM), time interacting with Stranger 1 and Stranger 2 (left), and Sociability Index (S.I.) for preference to interact with Stranger 2 (right). (E) Representative illustration of the Two-trial Place Recognition (2TPR) and Discrimination Index (D.I.) for time spent exploring the novel arm, compared to the familiar and start arms. (F) Representative illustration of the Novel Object Recognition (NOR) for short-term memory and Discrimination Index (D.I.) for time interacting with the new and familiar objects. (G) Representative illustration of the NOR for long-term memory and Discrimination Index (D.I.) for time interacting with new and familiar objects. (I) Spatial reference memory task of the Morris Water Maze (MWM) and respective Learning Curves for escape latency (left) and distance swam (right). (J) Reversal Learning task and time spent swimming in the old and new quadrants. Data plotted as mean ± SEM and analyzed using (A-G) Student’s t-test and (J-I) Two-way ANOVA test, being * *p* < .05, *** *p* < 0.001, and **** *p* < 0.0001. Control (Ctrl) mice are plotted in a black bar and individual values in black dots (n = 9-21) and dHIP:GFAP-mGluR5KO (KO) mice are plotted in a red bar and individual values in red dots (n = 10-23).

Following an emotional and social behavior assessment, mice were subjected to several behavioral paradigms to evaluate cognitive function. First, to assess place recognition memory, Control (n = 16) and dHIP:GFAP-mGluR5KO (n = 19) mice were tested in the 2TPR test, which specifically assesses place recognition memory (Figure 8E). The results obtained in the 2TPR test demonstrated that mice lacking astrocytic mGluR5 in the dorsal hippocampus presented an increased D.I. when compared to Control mice (Figure 8E, *p* = 0.02), suggesting an enhanced place recognition memory. Then, Control (n = 9-12) and dHIP:GFAP-mGluR5KO (n = 10-12) mice were tested in the NOR test to assess recognition memory (Figure 8F-G). In the short-term memory assessment (Figure 8F), our results showed that dHIP.GFAP-mGluR5KO and Control mice presented similar and positive D.I. when comparing the time interaction with the new and familiar object (Figure 8F). However, when tested for long-term memory (Figure 8G), dHIP:GFAP-mGluR5KO mice exhibited a decreased D.I. than Control mice (Figure 8G, *p* = 0.03), indicating that astrocytic mGluR5 plays a role in long-term and not short-term recognition memory.

Finally, to explore the involvement of the hippocampal astrocytic mGluR5 in spatial memory, Control (n = 15-19) and dHIP:GFAP-mGluR5KO (n = 22-23) mice were tested in the MWM to assess spatial reference memory and behavioral flexibility (Figure 8H-I). To assess spatial reference memory, mice had to learn the position of a hidden platform using visual cues displayed in the room. We observed that mice from both groups learned the position of the platform, and their performance increased similarly throughout the four days of testing (Figure 8H, *p* < 0.0001). On the fifth day, mice performed the reversal learning task, in which the platform was relocated to the opposite quadrant compared to the initial position. Our results revealed that Control and dHIP:GFAP-mGluR5KO mice learned the new position of the platform quadrant (Figure 8I). Post hoc analysis demonstrated that, as expected, Control mice spent more time in the new quadrant when compared to the old (Figure 8I, *p* < 0.0001). However, dHIP:GFAP-mGluR5KO mice spent similar time swimming in both quadrants, exhibiting lower swimming time in the new quadrant when compared to Control mice (Figure 8I, *p* = 0.0003). Thus, mice lacking astrocytic mGluR5 in the hippocampus presented impaired behavioral flexibility.

### Overexpression of mGluR5 in hippocampal astrocytes impaired short-term memory and improved behavioral flexibility

To further assess the regulatory capacity of mGluR5 signaling in astrocytes, we evaluated the consequences of mGluR5 upregulation on the processing of cognitive and emotional behaviors. To start, Control (n = 19) and dHIP:GFAP-mGluR5+ (n = 20) mice were tested in the L/D box test to assess anxious-like behavior (Figure 9A). Our results showed that mice from both genotypes spent a similar amount of time in the light zone (Figure 9A), indicating that overexpression of mGluR5 does not influence anxious-like behavior in mice. Then, mice (n_Control_ = 18 and n_dHIP:GFAP-mGluR5+_ = 17) were tested in the TST to evaluate learned helplessness, a type of depressive-like behavior (Figure 9B). We observed that dHIP:GFAP-mGluR5+ mice spent less time immobile when compared to Control mice (Figure 9B, *p* = 0.02), suggesting that overexpression of mGluR5 induces a sort of active coping in mice. Following emotional behavior assessment, Control (n = 7) and dHIP:GFAP-mGluR5+ (n = 11-12) mice were tested in the 3CST to evaluate social preference and memory (Figure 9C-D). First, mice performed the 3CST-SP (Figure 9C), and the results showed that dHIP-GFAP-mGluR5+ mice spent more time interacting with the empty cup (*p* = 0.03) and Stranger 1 (*p* = 0.001) when compared to Control mice (Figure 9C). Nevertheless, the S.I. was similar between mice from both groups, despite the increased interaction (Figure 9C), indicating intact sociability. Then, mice were tested in the 3CST-SM to evaluate social memory (Figure 9D). Here, we observed that mice with overexpression of mGluR5 in hippocampal astrocytes display increased time interacting with Stranger 1 when compared to Control mice (Figure 9D, *p* = 0.04), while time interacting with Stranger 2 was similar between groups (Figure 9D). Furthermore, dHIP:GFAP-mGluR5+ mice presented a lower S.I. compared to Control mice (Figure 9D, *p* = 0.02), indicating impaired social memory. To further investigate the effects of astrocytic mGluR5 overexpression on behavior, mice were tested in several behavioral paradigms to assess their cognitive abilities. Specifically, mice were tested in the 2TPR to evaluate place recognition memory, in the NOR to assess recognition memory, and in the MWM for spatial reference memory and behavioral flexibility, all of which are typically altered in neurobiological diseases. Firstly, Control (n = 16) and dHIP:GFAP-mGluR5+ (n = 19) mice performed the 2TPR test (Figure 9E). Our data revealed that dHIP:GFAP-mGluR5+ mice presented a lower D.I. compared to Control mice (Figure 9E, *p* = 0.007), indicating that mice with overexpression of mGluR5 spent less time exploring new locations compared to previously explored ones. In the NOR test, Control (n = 7-10) and dHIP:GFAP-mGluR5+ (n = 8-12) mice were tested for short-(Figure 9F) and long-term recognition memory (Figure 9G). Our data showed that in the short-term memory assessment, dHIP:GFAP-mGluR5+ mice presented a smaller and negative D.I. when compared to Control mice (Figure 9F, *p* = 0.001), which indicates that mice spent more time interacting with the familiar object rather than the new one. Interestingly, when mice were tested for long-term recognition memory, similar D.I. were observed between mice from both genotypes (Figure 9G). Thus, overexpression of mGluR5 in hippocampal astrocytes impairs short-term recognition memory without altering long-term recognition memory. Then, Control (n = 13-18) and dHIP:GFAP-mGluR5+ (n = 15-20) mice performed the MWM, which was divided into two tasks (Figure 9H-I). In the spatial reference task, where mice had to learn the position of a hidden platform, mice from both experimental groups presented a similar learning curve regarding time and distance swam to reach the platform throughout the four days of testing (Figure 9H, *p* < 0.0001). Lastly, Control and dHIP:GFAP-mGluR5+ mice were subjected to the reversal learning task, in which the platform was relocated to a different quadrant. Our results indicate that mice learned the new position of the platform (Figure 9I, *p* < 0.0001). Interestingly, dHIP:GFAP-mGluR5+ mice spent more time swimming in the new platform quadrant when compared to Control mice (Figure 9I, *p* = 0.05), which suggests enhanced behavioral flexibility.

**Figure 9.**
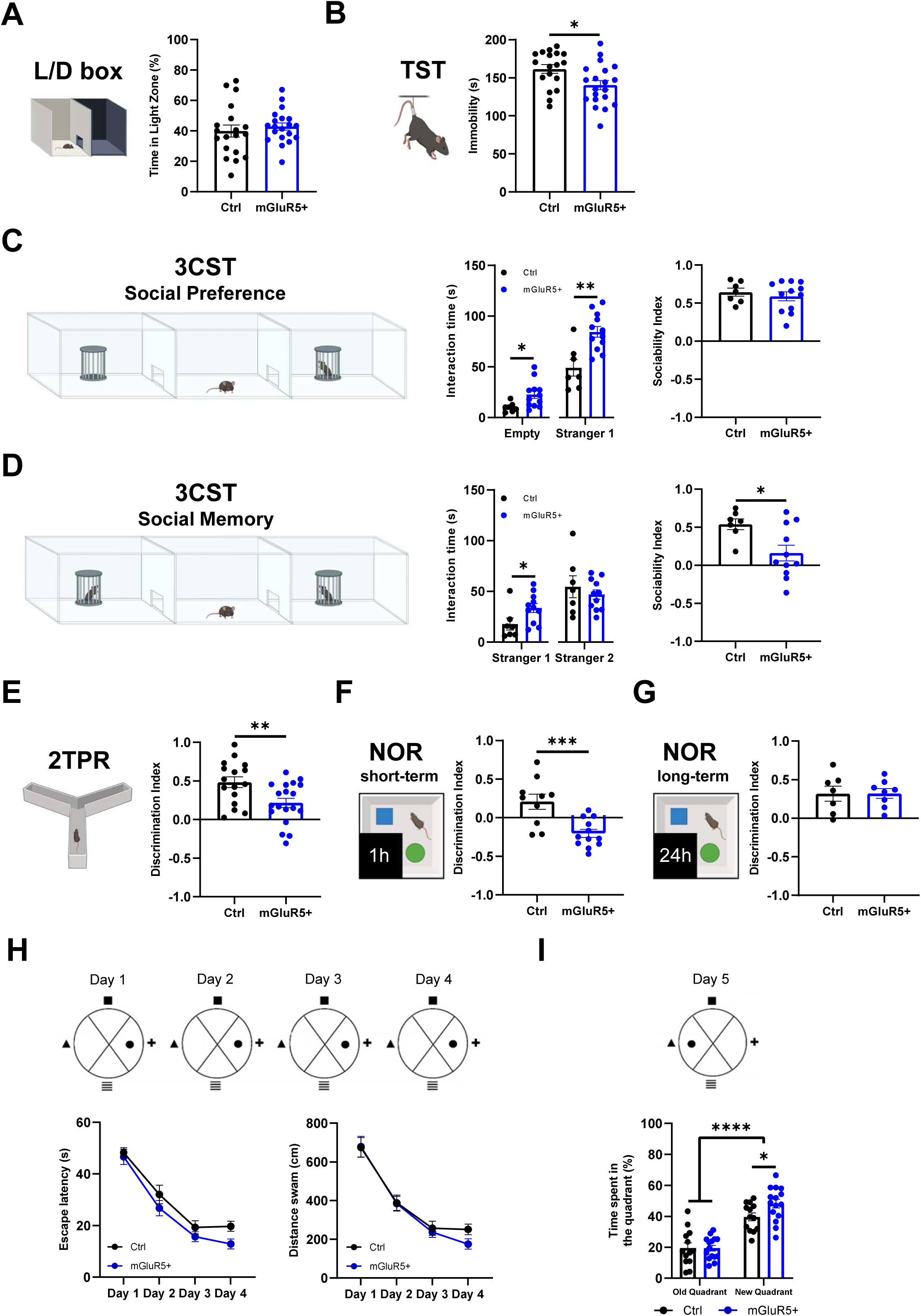
Overexpression of mGluR5 in hippocampal astrocytes increases active coping and induces social memory deficits. (A) Representative illustration of the Light/Dark (L/D) box test and percentage of time spent in the light zone of the L/D Box apparatus. (B) Representative Illustration of the Tail Suspension Test (TST) and immobility time in the TST. (C) Representative illustration of the 3 Chambers Social Test - Social Preference (3CST-SP), time interacting with the empty cup and Stranger 1 (left), and Sociability Index (S.I.) for preference to interact with Stranger 1 (right). (D) Representative illustration of the 3 Chambers Social Test - Social Memory (3CST-SM), time interacting with Stranger 1 and Stranger 2 (left), and Sociability Index (S.I.) for preference to interact with Stranger 2 (right). (E) Representative illustration of the Two-trial Place Recognition (2TPR) and Discrimination Index (D.I.) for time spent exploring the novel arm, compared to the familiar and start arms. (F) Representative illustration of the Novel Object Recognition (NOR) for short-term memory and Discrimination Index (D.I.) for time interacting with the new and familiar objects. (G) Representative illustration of the NOR for long-term memory and Discrimination Index (D.I.) for time interacting with new and familiar objects. (I) Spatial reference memory task of the Morris Water Maze (MWM) and respective Learning Curves for escape latency (left) and distance swam (right). (J) Reversal Learning task and time spent swimming in the old and new quadrants. Data plotted as mean ± SEM and analyzed using (A-G) Student’s t-test and (J-I) Two-way ANOVA test, being * *p* < .05, ** *p* < 0.01, *** *p* < 0.001, and **** *p* < 0.0001. Control (Ctrl) mice are plotted in a black bar and individual values in black dots (n = 7-19) and dHIP:GFAP-mGluR5+ (mGluR5+) mice are plotted in a blue bar and individual values in blue dots (n = 8-20).

## Discussion

### Astrocytes of adult mice express functional mGluR5

We conducted ex vivo Ca^2+^ imaging in acute hippocampal slices to assess astrocytic mGluR5 activity and subsequent deletion/overexpression. We observed that astrocytes from control mice displayed DHPG-evoked Ca^2+^ events using two different Ca^2+^ indicators, Fluor-4 and GCaMP6f. DHPG is an agonist of group I mGluRs, thereby activating mGluR1 and mGluR5. Given that an antagonist for mGluR1 was employed, we can confirm that DHPG-evoked Ca^2+^ events were mainly reliant on mGluR5 in astrocytes. Furthermore, the properties of the DHPG-evoked Ca^2+^ events were similar to those induced by agonists for different receptors (González-Arias et al., 2023), definitely confirming the existence of functional astrocytic mGluR5 in adulthood. Additionally, astrocytes lacking mGluR5 displayed reduced DHPG-evoked Ca^2+^ events, while astrocytes overexpressing mGluR5 displayed increased DHPG-evoked Ca^2+^ events. Notably, our results diverge from those of two studies that focus on astrocytic mGluR5 (Sun et al., 2013; Umpierre et al., 2019). The authors emphasize that astrocytes from adult mice do not respond with Ca^2+^ elevations in the presence of DHPG, concluding that astrocytic mGluR5 is not expressed in adult mice (Sun et al., 2013; Umpierre et al., 2019). However, a recent study indicated that astrocytes display mGluR5-dependent neuronal modulation in older mice, although this activity is reduced compared younger mice (Gómez-Gonzalo et al., 2017), supporting our findings. Moreover, in our experiments, we consistently observed Ca^2+^ responses, confirming the reliability of these findings. Panatier and Robitaille (2016) proposed that when recording mGluR5-dependent Ca^2+^ activity, one must consider the properties and localization of mGluR5. For instance, astrocytic mGluR5 has been shown to exhibit reduced desensitization compared to neuronal mGluR5 (Balázs et al., 1997). This reduced desensitization allows astrocytic mGluR5 to detect small increases in glutamate effectively. Finally, the heterogeneous expression of mGluR5 in different astrocyte populations of the hippocampus should be considered, as this region is known to include molecularly and structurally distinct populations (Batiuk et al., 2020; Viana et al., 2023). Overall, our findings enable us to conclude that astrocytic mGluR5 remains functional in mature astrocytes, and its activity can be modulated and studied.

### The biological relevance of astrocytic mGluR5 for neural network plasticity and behavioral processing

We generated a mouse model that allows the deletion of mGluR5 in astrocytes throughout the entire brain of adult mice. To our knowledge, this is the first study showing the relevance of this receptor in behavioral processing by the adult brain. Therefore, we conducted a comprehensive behavioral characterization of the ALDH1L1-mGluR5KO mouse model. We assessed various behavioral dimensions, including emotional regulation, sociability, learning, and memory, through complementary behavioral paradigms. We found that ALDH1L1-mGluR5KO mice exhibit an anxious-like phenotype and learned helplessness, accompanied by social deficits. Regarding cognition, the deletion of astrocytic mGluR5 impaired long-term recognition memory, while enhancing behavioral flexibility. Several studies have linked astrocytes to the various behavioral dimensions altered in the ALDH1L1-mGluR5KO mouse model (Oliveira et al., 2015; Nagai et al., 2021). For instance, research has demonstrated that the activation of astrocytes is crucial for processing novel and anxiogenic environments, playing a critical role in regulating anxious-like behaviors (Cho et al., 2022; Li et al., 2024). Likewise, astrocytes influence depressive-like behavior, particularly learned helplessness (Cao et al., 2013; Zhao et al., 2022), as well as sociability (Wang et al., 2021; González-Arias et al., 2023). Additionally, astrocytes have been directly linked to cognitive processing, influencing neuronal activity (Sardinha et al., 2017; Adamsky et al., 2018; Mederos et al., 2019; Navarrete et al., 2019). In the brain, two phenomena are associated with memory formation, retention, retrieval, and forgetting, specifically LTP and LTD. A balance between LTP and LTD is necessary for normal brain processing, affecting emotional regulation and cognitive function (Nabavi et al., 2014). Interestingly, astrocytes have been shown to facilitate both LTP (Henneberger et al., 2010; Abreu et al., 2023) and LTD (Navarrete et al., 2019; Pinto-Duarte et al., 2019), thereby influencing rodent behavior. Altogether, our findings led us to postulate that the activation of astrocytic mGluR5 by glutamate at the synaptic cleft triggers signaling cascades in astrocytes that might induce the release of gliotransmitters, including glutamate, D-serine, and ATP. These gliotransmitters will modulate synaptic plasticity, altering synaptic strength through LTP and LTD, which in turn modulate normal behavior in mice. To test this hypothesis, we need to understand how astrocytic mGluR5 modulates synaptic activity and identify the brain regions responsible for the observed behavioral outcomes. By merging our results with the available literature, two brain regions arise as crucial: the prefrontal cortex and the hippocampus. Within these regions, the hippocampus is one of the most extensively studied brain regions involved in all behavioral dimensions examined in this work. Thus, we decided to move forward focusing on astrocytic mGluR5 activity in the hippocampus. Then, we generated a mouse model lacking mGluR5 in astrocytes from the CA1 region of the dorsal hippocampus (dHIP:GFAP-mGluR5KO). After deletion of mGluR5, we observed that astrocytes exhibit diminished mGluR5-evoked Ca^2+^ elevations. Thus, confirming the deletion of the receptor. We then aimed to determine whether astrocytic mGluR5 activation modulates synaptic activity. We observed that astrocytic mGluR5 deletion enhanced basal transmission and, under high-frequency stimulation, abolished LTP. These synaptic alterations were translated into behavioral changes, specifically, dHIP:GFAP-mGluR5KO mice exhibit anxious-like behavior, social deficits, impaired long-term recognition memory and behavioral flexibility, but enhanced place recognition memory. Altogether, the findings in this work provide insight into the relevance of astrocytic mGluR5 for neuronal plasticity and hippocampal-dependent behavior, which facilitates a better understanding of the results obtained. Indeed, deletion of astrocytic mGluR5 in hippocampal astrocytes was sufficient to recapitulate the anxious-like phenotype, social deficit, and long-term recognition memory observed in the ALDH1L1-mGluR5KO mice. Interestingly, our results revealed that astrocytic mGluR5 has a regionally specific role in place recognition memory and behavioral flexibility. We observed that the deletion of astrocytic mGluR5 in the whole brain induced a marked anxious-like phenotype in mice. In contrast, deletion specifically in astrocytes of the dorsal hippocampus promoted a mild phenotypical change. Astrocytes in the hippocampus, specifically, in the DG region, have been linked to how rodents respond to anxiogenic environments (Cho et al., 2022; Li et al., 2024). However, it has also been demonstrated that hippocampal astrocytes play a pivotal role in the natural exploration of novel environments (Davis et al., 2004; Kemp and Manahan-Vaughan, 2004, 2008). Thus, activation of astrocytic mGluR5 in the CA1 region may be necessary for a normal environment assessment to regulate anxious-like behavior, implying that normal hippocampal network connectivity is essential to prevent an anxious-like state. More intriguingly, we observed enhanced behavioral flexibility when mGluR5 was deleted in astrocytes from the whole brain, while deletion specifically in the dorsal hippocampus impaired behavioral flexibility. Similarly, we observed that ALDH1L1-mGluR5KO mice exhibited intact place recognition memory, whereas deletion in the dorsal hippocampus induced an enhancement in this type of memory. Moreover, by overexpressing mGluR5 in astrocytes we observed the opposite effect in these specific behaviors. Thus, this indicates that modulation of these behaviors is directly dependent on astrocytic mGluR5 in the dorsal hippocampus. The discrepancy between the findings from our mouse models may be related to the extent of astrocytic manipulation, which could be compensating for these different behaviors. For instance, behavioral flexibility is classically associated with prefrontal cortex activity, whereas the hippocampus also contributes, namely by providing spatial cues in tasks such as the modified MWM (Nicholls et al., 2008; Mills et al., 2014). In our laboratory, we demonstrated that blocking the release of gliotransmission induces critical desynchronization of neuronal theta oscillations between the prefrontal cortex and the dorsal hippocampus (Sardinha et al., 2017). This desynchronization was strongly correlated with cognitive impairments in tasks dependent on this network (Sardinha et al., 2017). Thus, the activation of astrocytic mGluR5 in both regions may be crucial for normal behavioral flexibility, as evidenced by the enhanced behavioral flexibility observed when mGluR5 was deleted in both regions. However, when we directly ablated astrocytic mGluR5 in the dorsal hippocampus, an impaired behavioral flexibility was observed. Several studies have shown that LTD is crucial for normal behavioral flexibility (Nicholls et al., 2008; Dong et al., 2013; Mills et al., 2014), which was not assessed in this work. Here, we only evaluated LTP in dHIP:GFAP-mGluR5KO mice, which was abolished by mGluR5 deletion. Evaluating LTD may be required to fully understand the behavioral outcome obtained, as ALDH1L1-mGluR5KO mice display enhanced behavioral flexibility, which may indicate LTD facilitation.

### The impact of astrocytic mGluR5 overexpression in the dorsal hippocampus on synaptic plasticity and behavior

To study the consequences of astrocytic mGluR5 activation, and due to the lack of tools (e.g., pharmacology) to activate it specifically in astrocytes, we induced the overexpression of mGluR5 in astrocytes from the CA1 dorsal hippocampus of healthy mice. We observed that astrocytes overexpressing mGluR5 exhibit significant Ca^2+^ elevations triggered by DHPG, an agonist of group 1 mGluRs. These heightened astrocytic Ca^2+^ dynamics have been reported in contexts of neurobiological diseases, where they are often linked to glutamate toxicity (Casley et al., 2009; Grolla et al., 2013; Shrivastava et al., 2013; Umpierre et al., 2019; Danjo et al., 2022). Furthermore, to understand whether hippocampal synaptic plasticity was affected, we measured basal neuronal transmission and LTP. Surprisingly, we found that basal transmission was intact in dHIP:GFAP-mGluR5+ mice, indicating the existence of a resting state activation of astrocytic mGluR5 receptors during normal synaptic activity, where the added copies of this receptor were not significantly activated. However, under high-frequency stimulation, which elevates extracellular glutamate levels, the activation of mGluR5 facilitated LTD instead of LTP. In this work, we demonstrated that astrocytic mGluR5 plays a crucial role in facilitating LTP under healthy conditions. However, when overexpressed, astrocytic mGluR5 appears to shift synaptic plasticity towards LTD. A hypothesis for this shift from LTP to LTD might relate to the increased susceptibility of astrocytes to glutamate due to the overexpression of mGluR5. During periods of high synaptic activity, glutamate levels in the synaptic cleft increase. As astrocytes present a higher concentration of mGluR5, mGluR5-dependent activity will be amplified, potentially misleading astrocytes into sensing an abnormal rise in glutamate at the synaptic cleft. As demonstrated in this work, under basal conditions, astrocytic mGluR5 facilitates LTP. However, astrocytes overexpressing mGluR5 might redirect their synaptic plasticity component to facilitate LTD, thus reducing glutamate levels, a hypothesis that needs to be confirmed.

Our dHIP:GFAP-mGluR5+ mouse model displays a shift in synaptic plasticity, which might help clarify the behavioral readouts observed. These mice exhibited impaired social memory, short-term recognition memory, and place recognition memory, as well as enhanced behavioral flexibility. Interestingly, we showed that under basal conditions, astrocytic mGluR5 in the hippocampus facilitates LTP and is involved in anxious-like behavior, sociability, and long-term recognition memory—behavioral dimensions that seem intact in mice overexpressing mGluR5. Additionally, overexpressing mGluR5 resulted in opposite effects on immediate place recognition memory and behavioral flexibility. Thus, when mGluR5 is overexpressed in dorsal hippocampal astrocytes, these cells appear to shift their modulation toward more immediate/short-term memories and flexibility. Moreover, dHIP:GFAP-mGluR5+ mice showed reduced immobility in the TST. A decline in immobility could be associated with reduced behavioral despair. For instance, mice might shift their behavioral strategy to active coping, adapting better to stressful situations. Nonetheless, mice overexpressing mGluR5 also presented an increase in interaction time with other mice and objects, which could be associated with hyperactivity (Miyakawa et al., 2001; Zhuang et al., 2001). Although we observe a marked phenotype in dHIP:GFAP-mGluR5+ mice, their behavior does not fit the characteristics of a specific disease mouse model. To fully understand and correlate the obtained results, further experiments are necessary to elucidate why and how astrocytic mGluR5 overexpression modulates synaptic plasticity and consequently behavior. For instance, it would be interesting to explore how LTD is affected in the dHIP.GFAP-mGluR5+ mice, since high-frequency stimulation promotes synaptic depression. Moreover, it remains to determine the impact of astrocytic mGluR5 overexpression on slow inward currents (SICs), since it has been demonstrated that glutamate released by astrocytes upon mGluR5 activation binds specifically to extrasynaptic NMDARs, promoting SICs (Angulo et al., 2004; Fellin et al., 2004; Gómez-Gonzalo et al., 2017). These SICs were shown to modulate neuronal activity, being crucial for neuronal synchrony (Angulo et al., 2004; Fellin et al., 2004).

### Conclusion

Taken together, our results demonstrate that astrocytic mGluR5 is still biologically relevant in the adult brain. Here, we demonstrate that the ablation of astrocytic mGluR5 disrupts the normal astrocytic modulation of synaptic strength, which appears to have a particularly significant impact on hippocampal circuitry. Indeed, our findings suggest that mGluR5 activation in astrocytes is required for basal transmission and LTP facilitation. Furthermore, astrocytic mGluR5 is involved in emotional regulation, sociability, and cognitive function. Additionally, we demonstrated that activation of mGluR5 specifically in dorsal hippocampal astrocytes is required for these behaviors. Moreover, it also appears that the activation of astrocytic mGluR5 is region-specific, indicating that activation in different regions of the limbic system simultaneously or in response to another region may be crucial for normal behavioral processing. Notably, the deletion of astrocytic mGluR5 in the dorsal hippocampus was insufficient to recapitulate the learned helplessness and fear memory impairment observed, which may indicate the involvement of other brain regions, such as the prefrontal cortex and basolateral amygdala. Additionally, when mGluR5 was overexpressed in astrocytes, a shift in synaptic plasticity was observed, followed by impaired social memory, short-term recognition memory, and place recognition memory, as well as enhanced behavioral flexibility. Interestingly, a possible active coping mechanism was observed in these mice when they were exposed to stressful situations. Overall, this work provides evidence on the importance of astrocytic mGluR5 in adulthood, while bringing new insights into the involvement of this receptor in neurobiological diseases.

## Methods

### Animals

All procedures involving mice were performed in accordance with the guidelines for the welfare of laboratory animals, as described in Directive 2010/63/EU. In addition, they were approved by the National Authority for Animal Experimentation, DGAV (DGAV 023838/2019 and DGAV 34850/24-S) and the local Bioethics Committee of Cajal Institute, CSIC (2013/53/RD). Mice were group-housed in standard cages (3 to 6 mice per cage) with food and water *ad libitum*. The housing room was at 22 ± 1°C with controlled ventilation and was under a light/dark cycle of 12 h (light phase from 8 A.M. to 8 P.M.).

### Generation of a tamoxifen-inducible mouse model lacking astrocytic mGluR5

In this study three mouse lines, all in a C57BL/6J background, were used: mGluR5^flox/flox^ line, in which loxP sites flank the exon 7 of the mouse Grm5 gene, kindly provided by Dr. Anis Contractor [Northwestern University, Evanston, IL, USA, (Xu et al., 2009)]; ALDH1L1-Cre/ERT2 line [B6N.FVB-Tg(Aldh1l1-cre/ERT2)1Khakh/J RRID: IMSR_JAX:029655], in which the expression of tamoxifen-dependent Cre recombinase is mediated by the astrocyte-specific aldehyde dehydrogenase 1 family member L1 (ALDH1L1) promoter; ROSA26-stopflox-tdTomato cKI, which presents the Cre/loxP-dependent allele capable of driving high levels of cytosolic tdTomato expression, was gently provided by Professor Guoping Feng [Massachusetts Institute of Technology, Cambridge, MA, USA, (Arenkiel et al., 2011)]. The triple transgenic mouse line for knock-out (KO) of astrocytic mGluR5 was obtained as follows: First, the mGluR5^flox/flox^ line was crossed with the ALDH1L1-Cre/ERT2 line to generate the ALDH1L1-Cre/ERT2-mGluR5^flox/flox^ mouse. In parallel, the mGluR5^flox/flox^ line was also crossed with the ROSA26-stopflox-tdtomato line, to obtain mGluR5^flox/flox^-ROSA26-stopflox-tdtomato mouse. Then, the ALDH1L1-Cre/ERT2-mGluR5^flox/flox^ mouse was crossed with the mGluR5^flox/flox^-stopflox-tdtomato to generate the ALDH1L1-Cre/ERT2-mGluR5^flox/flox^-stopflox-tdTomato and mGluR5^flox/flox^-stopflox-tdTomato mice. All genotypes were identified by Polymerase Chain Reaction (PCR) analysis in DNA extracted from ear tissue. To identify Cre recombinase-expressing mice, the WT (forward, CTG TCC CTG TAT GCC TCT GG; reverse, AGA TGG AGA AAG GAC TAG GCT ACA) and mutant allele-specific primers (forward, CTT CAA CAG GTG CCT TCC A; reverse, GGC AAA CGG ACA GAA GCA) were used. For the flanked mGluR5 gene, the primers binding before and within the exon 7 (forward, AGA TGT CCC ACT TAC CTG ATG T; reverse, AGT TCC GTG TCT TTA TTC TTA GC) were used. Finally, primers to identify WT (forward, CAG TTG CTC TCC CAA AGT CG; reverse, GTT ATG TAA CGC GGA ACT CC) and tdTomato-expressing mice (forward, CAG TTG CTC TCC CAA AGT CG; reverse, TAG TCT AAC TCG CGA CAC TG) were used.

### In vivo tamoxifen injections

For the *in vivo* deletion of astrocytic mGluR5, male and female ALDH1L1-Cre/ERT2-mGluR5^flox/flox^-stopflox-tdTomato mice, 7 to 8 weeks old, were used. As controls, littermate mGluR5^flox/flox^-stopfloxtdTomato mice were used. Tamoxifen (Sigma-Aldrich, USA) was dissolved in corn oil (Sigma-Aldrich, USA) at a final concentration of 10 mg/mL and incubated overnight at 37 °C under agitation at 200–250 rpm. Mice were injected with 1 mg of tamoxifen twice a day for five consecutive days (daily dose: 2x 30-40 mg/kg body weight). After a 7-day resting period, the protocol was repeated (Mateus-Pinheiro et al., 2017). Four weeks after the last injection, the now ALDH1L1-CreERT2-mGluR5^del/del^-tdtomato (henceforth referred to as ALDH1L1-mGluR5KO) and mGluR5^flox/flox^-stopfloxtdTomato (henceforth referred to as Control) mice were used for experiments.

### Generation of a viral vector for overexpression of mGluR5

To induce overexpression of mGluR5 specifically in astrocytes, we decided to use an adeno-associated virus (AAV) packed with the Glial Fibrillary Acidic Protein (GFAP) 0.7 kilobase (kb) synthetic promoter derived from the human GFAP, a shorter version (3.576 kb) of the mGluR5 gene (Grm5), and a reporter protein. To validate if the shorter Grm5 would retain proper folding and functionality, an *in silico* confirmation was performed. Structural modeling and functional predictions confirmed that the shorter Grm5 was likely to maintain its natural conformation and function. Moreover, a FLAG tag was added to the N-terminal part of the protein, particularly after the Signal peptide and flanked by the addition GS dipeptide. This approach allowed us to generate the AAV5:GFAP(0.7)-FLAG-shortGrm5.

### Stereotaxic viral injection

Knockout of mGluR5 in astrocytes was achieved through an intracranial bilateral injection of AAV serotype 5 (AAV5) to induce the expression of Cre recombinase (AAV5.GFAP(0.7)-iCre-T2A-mCherry, titer: 3.7 x 10^11 genome copies (g.c.)/mL), purchased from Vector Biolabs (USA), into the dorsal hippocampus of mGluR5^flox/flox^ mice. Overexpression of mGluR5 in astrocytes was accomplished by the bilateral injection of a AAV5 that will induce the expression of a shorter mGluR5 gene (AAV5:GFAP(0.7)-FLAG-shortGrm5, titer: 1 x 10^9 vector genomes (v.g,)/μL), generated in collaboration with ViraVector (Coimbra, Portugal) into the dorsal hippocampus of C57BL/6J mice. Additionally, for *ex vivo* Ca^2+^ imaging, C57BL/6J mice were also injected with the AAV5-GFAP-cytoGCaMP6f (Addgene 52925; titer 1.3 x 10^13 g.c/ml), to induce expression of a genetically encoded Ca^2+^ indicator. Moreover, to control for the stereotaxic injection, an AAV5 to only induce expression of mCherry (AAV5:GFAP(0.7)-mCherry) was injected into the dorsal hippocampus of mGluR5^flox/flox^ (purchased in Vector Biolabs, titer 3.3×10^11 g.c./mL) and C57BL/6J (generated by ViraVector, titer 1 x 10^9 v.g./μL) mice.

Male mice, 8-12 weeks old, were injected intraperitoneally (i.p.) with buprenorphine (0.05 mg/Kg) for analgesia. Mice were anesthetized with sevoflurane (4% sevoflurane with O_2_) and placed on a heated plate in the stereotaxic apparatus. To avoid corneal drying, eyes were covered with Vaseline. The scalp was then incised along the rostral-caudal axis using a scalpel blade to expose the skull. Using a semi-automatic drill, two symmetrical holes were drilled into the skull at the coordinates determined from the bregma: 1.8 mm anteroposterior, ±1.3 mm mediolateral (Paxinos and Franklin, 2001). Mice were bilaterally injected directly into the dorsal hippocampus with 1 µL of the viral vector solution at a rate of 100 nL/min using a Hamilton syringe coupled to a 30-gauge needle (Hamilton, Switzerland), 1.3 mm below the brain surface (dorsoventral coordinate) (Paxinos and Franklin, 2001). At the end of each injection, the needle was pulled up 0.1 mm and left in place for 5 min to allow for proper viral vector diffusion. After the surgery, the incision was sutured and mice were subcutaneously injected with a multivitamin mixture (2-10 mL/kg; Duphalyte/Pfizer, USA). Mice were allowed to recover for four weeks before testing. The mGluR5fl/fl mice injected with AAV5:GFAP(0.7)-iCre-T2A-mCherry were expectedly mGluR5^del/del^ in hippocampal GFAP+ cells (henceforth referred to as dHIP:GFAP-mGluR5KO mice). Similarly, C57BL/6J mice injected with AAV5/GFAP(0.7)-FLAG-shortGrm5 were expectedly overexpressing mGluR5 in hippocampal GFAP+ cells (henceforth referred as dHIP:GFAP-mGluR5+ mice).

### Immunohistochemistry

ALDH1L1-mGluR5KO (n = 3) mice and respective controls (n = 3), were deeply anesthetized [ketamine (150 mg/kg) and medetomidine (0.3 mg/kg)] and transcardially perfused with 0.9% sodium chloride (NaCl), followed by perfusion with 4% paraformaldehyde (PFA) in PBS. Then, the brains were removed and kept in 4% PFA for 36 h. After this period, brains were transferred to a 30% sucrose solution until complete impregnation. Using a vibrating-blade microtome Leica VT 1000 S, free-floating 30 μm-thick slices were obtained. Free-floating tissue was first washed in PBS and then permeabilized with a 0.3% Triton X-100 (Sigma Aldrich, USA) in PBS (0.3% PBS-T) solution for 10 min. To reduce nonspecific binding, the tissue was incubated with 10% normal goat serum (NGS, Abcam) in 0.3% PBS-T blocking solution for 1 h at room temperature (RT). This blocking step was followed by overnight incubation at 4 °C with the primary antibodies rabbit anti-GFAP (1:200, Abcam, UK), rabbit anti-mGluR5 (1:200), rabbit polyclonal anti-NeuN (1:100; Cell Signalling, USA), rabbit polyclonal anti-iba1 (1:750; FUJIFILM Wako Chemicals, Hong Kong), rabbit polyclonal anti-S100β (1:200; Abcam, UK), rabbit polyclonal anti CC-1 (1:100, Millipore, USA), mouse anti-GFAP (1:200, Abcam, UK), and chicken anti-mCherry (1:500; HenBiotech, Portugal), in 2% NGS and 0.3% PBS-T. On the next day, the slices were washed in 0.3% PBS-T and incubated with the respective species-specific secondary antibody: goat anti-rabbit Alexa Fluor® 488 (1:1000; Thermo Fischer Scientific, USA), goat anti-mouse Alexa Fluor® 594 (1:1000; Thermo Fischer Scientific, USA), goat anti-chicken Alexa Fluor®594 (1:1000; Invitrogen, TermoFisher Scientific, USA), or goat anti-mouse®647 (1:1000; Thermo Fischer Scientific, USA), in a 0.3% PBS-T solution with 2% NGS, for 1 h at RT. After this period, the slices were washed with 0.3% PBS-T, and the nucleic acids were labeled with 4′,6-diamidino-2-phenylindole (DAPI) (1:1000, Invitrogen, USA) for 10 min at RT. Slices were washed with PBS and then mounted using Immu-MountTM (ThermoFisher Scientific, USA). All procedures performed on this day were conducted in the dark. Images for cellular specificity were acquired using a confocal microscope (FV1000, Olympus, Japan) with a 60x objective, a 1 µm z-step, and a resolution of 640×640 pixels. The images for fluorescence quantification were acquired in a widefield inverted microscope with 20x magnification (Olympus, Japan) in four different fluorescence channels (TRITC, FITC, and DAPI) with the same acquisition parameters.

### Fluorescence quantification

Fluorescence quantification of astrocytic mGluR5 was conducted using the Image J software (https://fiji.sc/). Astrocytes for mGluR5 fluorescence quantification were selected based on the presence of a DAPI-stained nucleus, tdTomato/GFAP-marked processes radiating from the nucleus. The regions of interest (ROI) were manually defined using the brush tool, surrounding the astrocytic territory delimited by the union of the ends of the tdTomato-labelled processes. Fluorescent channels were then separated (ImageJ: Image > Color > Split Channels), generating individual images for tdTomato, mGluR5, and DAPI. The predefined ROI was applied to the mGluR5 image, and areas outside the ROI were cleared (Edit > Clear Outside) to isolate the fluorescence, and mean fluorescence intensity within each ROI was measured (Ctrl + M). The mean intensity of a background area was also measured in the same image to correct the fluorescence signal measurement. The final mGluR5 fluorescence intensity was obtained by subtracting the mean intensity value for the background from the mean intensity of each ROI.

### Acute hippocampal slice preparation for ex vivo recordings

Mice (12 to 14 weeks old) were anesthetized with sodium pentobarbital (50–90 mg/kg) and perfused with NMDG-HEPES solution at 4 °C, to improve slice viability, followed by quick and careful brain removal. Subsequently, the brain was kept in ice-cold NMDG-HEPES gassed with carbogen (95% O_2_/5% CO_2_, pH = 7.3) and sectioned in a vibratome (Leica Vibratome VT1200S, Germany) into 350 µm-thick acute hippocampal slices. Then, the slices were placed in NMGD-HEPES at 32-34 °C for 10 min, followed by incubation for at least 1 h at RT, in artificial cerebrospinal fluid (aCSF) continuously gassed with carbogen (95% O_2_/5% CO_2_, pH = 7.3) until recordings were made. The NMDG-HEPES contained the following [in mM]: NMDG 92, KCl 2.5, NaH_2_PO_4_ 1.2, NaHCO_3_ 30, HEPES 20, glucose 25, thiourea 2, Na-ascorbate 5, Na-pyruvate 3, CaCl_2_·2H_2_O 0.5, and MgSO_4_·7H_2_O 10 gassed with carbogen (95% O2/5% CO2, pH = 7.3). The aCSF contained the following [in mM]: NaCl 124, KCl 2.69, KH_2_PO_4_ 1.25, MgSO_4_ 2, NaHCO_3_ 26, CaCl_2_ 2, and glucose 10, and gassed with carbogen (95% O_2_/5% CO_2_, pH = 7.3).

### Ex vivo astrocytic calcium imaging and analysis

For *ex vivo* Ca^2+^ imaging in acute hippocampal slices, two different Ca^2+^ indicators were used. The fluorescent Ca^2+^ indicator dye Fluor-4-AM was used to detect Ca^2+^ dynamics in astrocytes of dHIP:GFAP-mGluR5KO (n = 3) and respective Control (n = 2) mice. Acute hippocampal slices were incubated with Fluo-4 AM (1 μl of 2 mM dye at a final concentration of 2-10 μM, Invitrogen) dissolved in 0.02 % Pluronic® F-127 (Merck) and 0.04 % dimethyl sulfoxide (DMSO, Sigma-Aldric) for 15-20 min at RT, and Ca^2+^ dynamics analysis was restricted to the soma. Alternatively, the genetically encoded Ca^2+^ indicator GCaMP6f was used, delivered by the injection of the AAV5-GFAP-cytoGCaMP6f into dHIP:GFAP-mGluR5KO+ (n = 3) and Control (n = 3) mice., the

For recordings, slices were transferred to an immersion recording chamber superfused at 2 mL/min with gassed aCSF in the presence of tetrodotoxin (1 µM, Tocris), picrotoxin (50 µM, Sigma), AM251 (2 µM, Tocris), MRS 2179 tetrasodium salt (10 µM, Tocris), CGP 55845 hydrochloride (5 µM, Tocris), and LY367385 (100 µM, Tocris) to isolate mGluR5-dependent activity. Slices were left to rest in the chamber for 15 min before the recordings began. Imaging of astrocytic Ca^2+^ dynamics was performed using a CCD camera (ORCA-235, Hamamatsu, Japan) attached to the microscope in the CA1 *stratum oriens* and *stratum radiatum* of the dorsal hippocampus. Cells were illuminated for 100–200 ms at 490 nm using an LED system (CoolLED pE-100), which was controlled and synchronized with the CCD camera by NIS Elements software (Nikon, Japan). Images were acquired at 1 Hz for 2 min. Evoked Ca^2+^ responses were measured by recording baseline activity for 30 s, followed by local application of (S)-3,5-Dihydroxyphenylglycine (DHPG, 1 mM) by pressure pulses through a micropipette (5 s, 1 bar; Picospritzer II, Parker Hannifin, Mayfield Heights, OH, USA). Agonist-evoked activity that was restricted to 60 s upon stimulation. For dHIP:GFAP-mGluR5+ and respective Control mice, spontaneous Ca^2+^ dynamics were also recorded for 2 min.

To analyze the recorded Ca^2+^ dynamics, ROIs were manually selected, and all pixels were averaged to obtain a single time course F[t] per ROI using ImageJ software. Further processing was performed using custom-written software in MATLAB (MATLAB R2021a; MathWorks, Natick, MA), as described in González-Arias *et al*., 2023. Artifacts produced by mechanical movement were removed from the analysis. Signals were then low-pass filtered using a Chebyshev II filter, followed by adjustment for photobleaching and calculation of ΔF/F_0_ for each ROI. Calcium events were considered when ΔF/F_0_ > 2-3 times the noise variance and had at least > 3% of relative change (0.03). For each ROI, frequency, amplitude, area under the curve, and duration were obtained.

### Extracellular recordings of field excitatory postsynaptic potentials

Acute hippocampal slices from dHIP:GFAP-mGluR5KO (n = 4), dHIP:GFAP-mGluR5+ (n = 5), and respective Control (n = 3) were transferred to a recording chamber superfused, at a flow of 2 mL/min, with aCSF with picrotoxin 50 µM gassed with carbogen and kept at 37 °C. Before extracellular recordings of the field excitatory postsynaptic potential (fEPSP), slices were allowed to rest in the chamber for 15 min, to allow picrotoxin to act. Recording extracellular fEPSPs was performed by placing a recording electrode in the CA1 *stratum radiatum* of the dorsal hippocampus, and stimulation was applied through a bipolar tungsten wire electrode (WPI TST33A05KT) positioned in the Schaffer collaterals. The fEPSPs were evoked by the stimulation (20 µs pulse duration) of the Schaffer collateral fibers, using a DS3 Isolated Current Stimulator (Digitimer, UK). Stimulation intensity was adjusted to obtain a fEPSP slope corresponding to 50% of the maximum evoked fEPSP slope. Basal fEPSPs were recorded using constant stimulus intensity and frequency, and fEPSP slope stability was monitored for at least 10 min before recording. Twenty consecutive fEPSPs responses were acquired at 0.1 Hz using a Model 3000 AC/DC differential amplifier (A-M Systems, USA), digitized with a Digidata 132 Axon Instruments 16-Bit Acquisition system (Axon Instruments, USA). To remove noise in the fEPSP recordings, a HumBug 50/60 Hz Noise Eliminator (Digitimer, UK) was used. Recorded fEPSPs were quantified by measuring the slope (duration 0.4ms) of the averaged 20 fEPSPs per min in the Clampfit 11.3 software (Molecular Devices, USA). For each experimental group, two to three acute hippocampal slices were used to generate Input/Output curves and induce Long-term Potentiation (LTP).

### Input/Output curves

To measure basal synaptic transmission, after obtaining a stable fEPSP baseline activity in the CA1 *stratum radiatum* of the dorsal hippocampus, the stimulus intensity was decreased to 0 µA and then increased stepwise until 1300 µA. For each stimulation intensity, five consecutive fEPSPs were recorded and averaged to measure each fEPSP slope.

### Long-term potentiation induction

Long-term potentiation (LTP) was induced in acute hippocampal slices after recording a stable fEPSP baseline for at least 15 min. To induce LTP, a theta burst stimulation protocol consisting of 4 bursts of 100 Hz each, separated by 200 ms, was repeated 4 times with a 20 s interval. The magnitude of LTP was quantified as the change ratio of the fEPSP slope in the last 10 min compared to the average fEPSP slope.

### Behavioral testing

Independent sets of ALDH1L1-mGluR5KO male (n = 9-30) and female (n = 9-25) mice and respective littermate controls (n_males_ = 9-28 and n_females_ = 8-21) were tested to dissect the involvement of astrocytic mGluR5 in behavior. Furthermore, two independent sets of dHIP:GFAP-mGluR5KO (n = 10 - 23), dHIP:GFAP-mGluR5+ (n = 8 - 20), and respective littermate controls (n = 9 - 21 and n = 7 - 19) were tested to dissect the involvement of astrocytic mGluR5 in behavior dependent on the hippocampus. Mice were handled daily for 5 min over the week preceding testing. On the test day, mice were habituated to the testing room for 30 min before behavioral evaluation. All behavior tests were conducted during the light phase.

### Light/Dark Box

The Light/Dark (LD) box test was used to assess anxious-like behavior in rodents. This test is based on the natural aversion of rodents to brighter areas and their propensity to explore an arena in response to mild stressors such as a novel environment and light (Bourin and Hascoët, 2003). The LD box apparatus consists of an arena (43.2 x 43.2 x 30.5 cm) equally divided into light and dark compartments, connected by an opening (Med Associates Inc, Vermont, USA). Mice were placed in the middle of the illuminated compartment, facing toward the dark area, and allowed to explore the arena for 10 min. Data was analyzed manually, and time in the light compartment was used as an index of anxious-like behavior.

### Tail Suspension Test

The Tail Suspension Test (TST) was used to evaluate learned helplessness since rodents tend to develop an immobile position when placed in an unavoidable stressful situation (Can et al., 2012b). Mice were suspended by their tails, on the edge of a shelf, using adhesive tape, 80 cm above the floor, for 6 min, and behavioral activity was recorded using a video camera. EthoVision XT 13.0 (Noldus Information Technology, Netherlands) was used to obtain, for each mouse, the period of immobility during the final 4 min of the test, as the mice had to learn that there was no possible escape.

### Forced Swim Test

The Forced Swim Test (FST) was performed to measure learned helplessness in mice (Can et al., 2012a), based on the same principle as the TST. In the FST, mice were placed in a transparent glass cylinder (15 cm in diameter) filled with water at 23 °C. The test had a duration of 5 min, and the time spent immobile in the last 3 min was used as a measure of learned helplessness. The test was video recorded and analyzed using the ANY-maze Video Tracking System version 7.35 (Stoelting Europe, Ireland).

### Three-Chamber Sociability and Social Memory Test

The Three-Chamber Social test (3CST) was performed to evaluate social preference and memory of mice (Kaidanovich-Beilin et al., 2011). The test was performed in an open arena (75 L x 50 W x 25 H cm) subdivided into three equal and connected chambers with a wire cup in the left and right chambers. The test had a total duration of 30 min and was subdivided into three consecutive phases of 10 min each. In the first phase, the habituation phase, mice were placed in the middle chamber and allowed to explore the arena for 10 min. In phase two, the social preference test (3CST-SP), a strange mouse (Stranger 1) was placed under one of the wire cups while the other remained empty, and the test mouse could explore the arena for 10 min. Social preference was evaluated by measuring the time mice spent interacting with the test mouse in the wire cup compared to the empty wire cup. Finally, in the third phase, social memory test (3CST-SM), a new stranger mouse (Stranger 2) was placed in the previous empty wire cup, and the mouse was allowed to explore the arena for another 10 min. Social memory was evaluated by comparing the time the test mouse spent interacting with the unknown mouse (Stranger 2) and the familiar mouse (Stranger 1). To control for the interaction time, in seconds, of each mouse, a sociability index (S.I.) was calculated using the following equations (1) and (2):

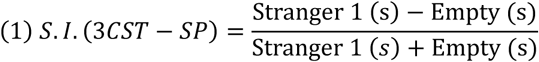

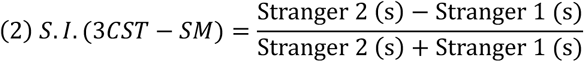

The Stranger 1 and 2 mice, age-matched to the test mouse, were exchanged frequently and placed in alternate positions in the arena, to disclose any place preference from the test mouse. The test was performed under dim light conditions, and the arena was cleaned with 10% ethanol between mice.

### Two-Trial Place Recognition

The Two-Trial Place Recognition (2TPR) test was used to assess place recognition memory and is based on the native propensity of rodents to explore novel locations (Guerra-Gomes et al., 2018). The 2TPR was performed in a Y-shaped arena composed of three equal arms (33.2 L x 7 W x 15 H cm) designated as Start (S), Familiar (F), or Novel (N) arms, made of white Plexiglas. At the end of each arm, a visual cue was added to allow mice to recognize each arm based on their spatial recognition and navigation. In the first trial, mice were placed individually in the S arm and allowed to explore the S and F arms for 5 min. Immediately after, all arms were available, and mice were placed again in the S arm and left to explore the arena for 2 min. The test was performed under dim light conditions, and the maze was cleaned with 10% ethanol between phases and subjects. All trials were recorded using a video camera and further analyzed using EthoVision XT 13.0 (Noldus Information Technology, Netherlands) software. A Discrimination Index (D.I.) for time spent in each arm, in seconds, was calculated using the following equation (3):

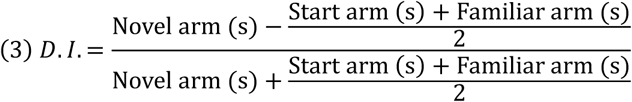

### Novel Object Recognition

The Novel Object Recognition (NOR) test was performed to assess recognition memory, based on the natural tendency for mice to prefer to explore novelty (Lueptow, 2017). The test was performed in an L-shaped apparatus (44 cm and 34 cm long for external and internal walls, respectively, 10 cm wide and 44 cm high walls). On the first day, mice were allowed to explore the empty apparatus for 10 min. After 24 h, familiarization phase, mice were placed for 10 min in the same apparatus with two similar objects (familiar objects) placed on each extremity of the apparatus. Then, in the testing phase, one of the objects was changed to a different object (novel object), and mice were placed in the apparatus for 5 min, with a retention interval of 1 h, to assess short-term memory or 24h to assess long-term memory. All trials were conducted under dim light conditions, and the apparatus and objects were cleaned with 10% ethanol between mice to eliminate odor cues. Each trial was recorded with a video camera and analyzed manually. Total object exploration time, in seconds, considered when mice smell, touch, or climb the objects, was extracted to calculate the discrimination index (D.I.) according to the following equation (4):

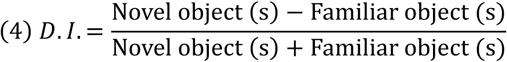

### Morris Water Maze

The Morris Water Maze (MWM) was performed to evaluate spatial reference memory and behavioral flexibility (Morris, 1984). The MWM apparatus consists of a dark circular pool (106 cm diameter) filled with water at 23°C, divided into four imaginary quadrants, which were associated with extrinsic visual cues (cross, lines, triangle, and square). In one of the imaginary quadrants, a circular escape platform (11 cm diameter, 26 cm height) was placed 1 cm below the water surface. To hide the platform and increase contrast for mouse detection, non-toxic titanium dioxide (Sigma-Aldrich; 250 mg/L) was added to the water. The MWM test was conducted under dim light conditions.

The Reference Memory task is based on the capacity of mice to learn the position of the escape platform, which remained in the same quadrant for four consecutive days. Each day, mice performed four trials with a maximum duration of 60 s. In each trial, mice were placed in the pool facing the pool wall and oriented to each of the visual cues in a random order. The trial ended when mice reached the hidden platform or failed to find it within 60 s. Whenever mice were unable to complete a trial, mice were guided to the platform and allowed to stay on it for 20 s. On the fifth day, a probe trial was conducted, during which the hidden platform was removed from the pool. Mice were allowed to explore the pool for 30 s, and the time spent in the previously established platform quadrant was then measured. If the mice learned the position of the escape platform, they would spend more time in the quadrant where the platform was hidden before. Escape latencies and distance swam were recorded and further analyzed using EthoVision XT 13.0 (Noldus Information Technology, Netherlands). Swimming patterns were also extracted using Ethovision software and then classified according to Graziano et al. (2003) and as described by Sardinha et al. (2017). To assess behavioral flexibility, a reversal learning task was conducted on the fifth day of the protocol. In this task, the platform was placed in the opposite quadrant as compared to its initial location during the reference memory task. Mice had four trials of 60 s to find the platform in the new quadrant. In the reversal learning task, the time spent and distance swam in the old and new platform quadrants were measured to evaluate their ability to learn a new rule. All trials were recorded using a video camera and analyzed using EthoVision XT 13.0 (Noldus Information Technology, Netherlands).

### Statistical Analysis

Statistical analysis was performed using the GraphPad Prism software, version 8.4.2 for Windows (GraphPad software, La Jolla, CA, USA). All data passed the Kolmogorov-Smirnov normality test for normal distribution. Statistical outliers were identified using Grubbs’ outlier test and excluded from the analysis. Moreover, mice that did not fulfill the behavioral test criteria were removed from the analysis, for example, did not interact with the mouse/objects, or did not learn how to perform the test. Comparisons between two groups were performed using an independent Student’s t-test. Analysis of variances (ANOVA) followed by Sidak post hoc analysis was used for comparisons between different variables. Statistically significant differences were considered when p < 0.05. Effect size measures (Cohen’s d for student’s t-test, partial eta squared (ɳ_p2_) for Two-way ANOVA were calculated. The values are presented as mean ± standard error of the mean (SEM).

## Acknowledgements and funding information

The authors are grateful to Professor Anis Contractor and Professor Guoping Feng for sharing the mouse lines. The authors acknowledge the Foundation for Science and Technology (FCT) fellowships to JFV, SB, DSA and JLM; CEECINST/00018/2021 Grant to JFO; grant from “la Caixa” Foundation (LCF/PR/HR21/52410024) to JFO; all authors were funded by FCT projects UIDB/50026/2020 (DOI 10.54499/UIDB/50026/2020), UIDP/50026/2020 (DOI 10.54499/UIDP/50026/2020), and LA/P/0050/2020 (DOI 10.54499/LA/P/0050/2020). ICVS Scientific Microscopy Platform, member of the national infrastructure PPBI - Portuguese Platform of Bioimaging (PPBI-POCI-01-0145-FEDER-022122; by National funds, through the Foundation for Science and Technology (FCT) - project UIDB/50026/2020. Grant PID2022-142617NBI00 to GP supported by Ministerio de Ciencia, Innovación y Universidades, Spain, MCIN/AEI/10.13039/501100011033 and “ERDF A way of making Europe”.

## Author contributions

JFV, GP and JFO designed the project, supervised the data analysis, and wrote the manuscript. JFV, JDD, CGA, LSA, AV, DSA, SB, and JLM performed the experiments and analyzed the results. LPA and RJN designed and generated the viral vector for overexpression used in the work. GP and JFO contributed to the intellectual content and critical manuscript review.

## References

Abreu DS, Gomes JI, Ribeiro FF, Diógenes MJ, Sebastião AM, Vaz SH. 2023. Astrocytes control hippocampal synaptic plasticity through the vesicular-dependent release of D-serine. Front Cell Neurosci 17:1282841.

Adamsky A, Kol A, Kreisel T, Doron A, Ozeri-Engelhard N, Melcer T, Refaeli R, Horn H, Regev L, Groysman M, London M, Goshen I. 2018. Astrocytic Activation Generates De Novo Neuronal Potentiation and Memory Enhancement. Cell 174:59–71.e14.

Angulo MC, Kozlov AS, Charpak S, Audinat E. 2004. Glutamate released from glial cells synchronizes neuronal activity in the hippocampus. J Neurosci 24:6920–6927.

Araque A, Carmignoto G, Haydon PG, Oliet SHR, Robitaille R, Volterra A. 2014. Gliotransmitters Travel in Time and Space. Neuron 81:728–739.

Arenkiel BR, Hasegawa H, Yi JJ, Larsen RS, Wallace ML, Philpot BD, Wang F, Ehlers MD. 2011. Activity-Induced Remodeling of Olfactory Bulb Microcircuits Revealed by Monosynaptic Tracing. PLOS ONE 6:e29423.

Batiuk MY, Martirosyan A, Wahis J, de Vin F, Marneffe C, Kusserow C, Koeppen J, Viana JF, Oliveira JF, Voet T, Ponting CP, Belgard TG, Holt MG. 2020. Identification of region-specific astrocyte subtypes at single cell resolution. Nat Commun 11:1220.

Bourin M, Hascoët M. 2003. The mouse light/dark box test. Eur J Pharmacol 463:55–65.

Can A, Dao DT, Arad M, Terrillion CE, Piantadosi SC, Gould TD. 2012a. The Mouse Forced Swim Test. J Vis Exp:3638.

Can A, Dao DT, Terrillion CE, Piantadosi SC, Bhat S, Gould TD. 2012b. The tail suspension test. J Vis Exp:e3769.

Cao X, Li L-P, Wang Q, Wu Q, Hu H-H, Zhang M, Fang Y-Y, Zhang J, Li S-J, Xiong W-C, Yan H-C, Gao Y-B, Liu J-H, Li X-W, Sun L-R, Zeng Y-N, Zhu X-H, Gao T-M. 2013. Astrocyte-derived ATP modulates depressive-like behaviors. Nat Med 19:773–777.

Casley CS, Lakics V, Lee H, Broad LM, Day TA, Cluett T, Smith MA, O’Neill MJ, Kingston AE. 2009. Up-regulation of astrocyte metabotropic glutamate receptor 5 by amyloid-β peptide. Brain Research 1260:65–75.

Cho W-H, Noh K, Lee BH, Barcelon E, Jun SB, Park HY, Lee SJ. 2022. Hippocampal astrocytes modulate anxiety-like behavior. Nat Commun 13:6536.

Danjo Y, Shigetomi E, Hirayama YJ, Kobayashi K, Ishikawa T, Fukazawa Y, Shibata K, Takanashi K, Parajuli B, Shinozaki Y, Kim SK, Nabekura J, Koizumi S. 2022. Transient astrocytic mGluR5 expression drives synaptic plasticity and subsequent chronic pain in mice. J Exp Med 219:e20210989.

Davis CD, Jones FL, Derrick BE. 2004. Novel Environments Enhance the Induction and Maintenance of Long-Term Potentiation in the Dentate Gyrus. J Neurosci 24:6497–6506.

Dias JD, Viana JF, Alves LS, Veiga A, Matos B, Machado JL, Oliveira JF. 2025. AstroWars: the return of the astrocytic metabotropic glutamate receptor 5. The Journal of Physiology [Internet] n/a. Available from: https://onlinelibrary.wiley.com/doi/abs/10.1113/JP288403

Fellin T, Pascual O, Gobbo S, Pozzan T, Haydon PG, Carmignoto G. 2004. Neuronal synchrony mediated by astrocytic glutamate through activation of extrasynaptic NMDA receptors. Neuron 43:729–743.

Ferrando RE, Newton K, Chu F, Webster JD, French DM. 2015. Immunohistochemical Detection of FLAG-Tagged Endogenous Proteins in Knock-In Mice. J Histochem Cytochem 63:244–255.

Gasiorowska A, Wydrych M, Drapich P, Zadrozny M, Steczkowska M, Niewiadomski W, Niewiadomska G. 2021. The Biology and Pathobiology of Glutamatergic, Cholinergic, and Dopaminergic Signaling in the Aging Brain. Front Aging Neurosci 13:654931.

Gómez-Gonzalo M, Martin-Fernandez M, Martínez-Murillo R, Mederos S, Hernández-Vivanco A, Jamison S, Fernandez AP, Serrano J, Calero P, Futch HS, Corpas R, Sanfeliu C, Perea G, Araque A. 2017. Neuron–astrocyte signaling is preserved in the aging brain. Glia 65:569–580.

González-Arias C, Sánchez-Ruiz A, Esparza J, Sánchez-Puelles C, Arancibia L, Ramírez-Franco J, Gobbo D, Kirchhoff F, Perea G. 2023. Dysfunctional serotonergic neuron-astrocyte signaling in depressive-like states. Mol Psychiatry 28:3856–3873.

Graziano A, Petrosini L, Bartoletti A. 2003. Automatic recognition of explorative strategies in the Morris water maze. J Neurosci Methods 130:33–44.

Grolla AA, Sim JA, Lim D, Rodriguez JJ, Genazzani AA, Verkhratsky A. 2013. Amyloid-β and Alzheimer’s disease type pathology differentially affects the calcium signalling toolkit in astrocytes from different brain regions. Cell Death Dis 4:e623.

Guerra-Gomes S, Viana JF, Nascimento DSM, Correia JS, Sardinha VM, Caetano I, Sousa N, Pinto L, Oliveira JF. 2018. The Role of Astrocytic Calcium Signaling in the Aged Prefrontal Cortex. Front Cell Neurosci 12:379.

Henneberger C, Papouin T, Oliet SHR, Rusakov DA. 2010. Long-term potentiation depends on release of D-serine from astrocytes. Nature 463:232–236.

Kaidanovich-Beilin O, Lipina T, Vukobradovic I, Roder J, Woodgett JR. 2011. Assessment of social interaction behaviors. J Vis Exp:2473.

Kemp A, Manahan-Vaughan D. 2004. Hippocampal long-term depression and long-term potentiation encode different aspects of novelty acquisition. Proc Natl Acad Sci U S A 101:8192–8197.

Kemp A, Manahan-Vaughan D. 2008. The Hippocampal CA1 Region and Dentate Gyrus Differentiate between Environmental and Spatial Feature Encoding through Long-Term Depression. Cerebral Cortex 18:968–977.

Li J-T, Jin S-Y, Hu J, Xu R-X, Xu J-N, Li Z-M, Wang M-L, Fu Y-W, Liao S-H, Li X-W, Chen Y-H, Gao T-M, Yang J-M. 2024. Astrocytes in the Ventral Hippocampus Bidirectionally Regulate Innate and Stress-Induced Anxiety-Like Behaviors in Male Mice. Advanced Science 11:2400354.

Lueptow LM. 2017. Novel Object Recognition Test for the Investigation of Learning and Memory in Mice. J Vis Exp:55718.

Mateus-Pinheiro A, Alves ND, Patrício P, Machado-Santos AR, Loureiro-Campos E, Silva JM, Sardinha VM, Reis J, Schorle H, Oliveira JF, Ninkovic J, Sousa N, Pinto L. 2017. AP2γ controls adult hippocampal neurogenesis and modulates cognitive, but not anxiety or depressive-like behavior. Mol Psychiatry 22:1725–1734.

Mederos S, Hernández-Vivanco A, Ramírez-Franco J, Martín-Fernández M, Navarrete M, Yang A, Boyden ES, Perea G. 2019. Melanopsin for precise optogenetic activation of astrocyte-neuron networks. Glia 67:915–934.

Mills F, Bartlett TE, Dissing-Olesen L, Wisniewska MB, Kuznicki J, Macvicar BA, Wang YT, Bamji SX. 2014. Cognitive flexibility and long-term depression (LTD) are impaired following β-catenin stabilization in vivo. Proceedings of the National Academy of Sciences 111:8631– 8636.

Miyakawa T, Yamada M, Duttaroy A, Wess J. 2001. Hyperactivity and Intact Hippocampus-Dependent Learning in Mice Lacking the M1 Muscarinic Acetylcholine Receptor. J Neurosci 21:5239–5250.

Morris R. 1984. Developments of a water-maze procedure for studying spatial learning in the rat. J Neurosci Methods 11:47–60.

Nabavi S, Fox R, Proulx CD, Lin JY, Tsien RY, Malinow R. 2014. Engineering a memory with LTD and LTP. Nature 511:348–352.

Nagai J, Yu X, Papouin T, Cheong E, Freeman MR, Monk KR, Hastings MH, Haydon PG, Rowitch D, Shaham S, Khakh BS. 2021. Behaviorally consequential astrocytic regulation of neural circuits. Neuron 109:576–596.

Navarrete M, Cuartero MI, Palenzuela R, Draffin JE, Konomi A, Serra I, Colié S, Castaño-Castaño S, Hasan MT, Nebreda ÁR, Esteban JA. 2019. Astrocytic p38α MAPK drives NMDA receptor-dependent long-term depression and modulates long-term memory. Nature Communications 10:2968.

Nicholls RE, Alarcon JM, Malleret G, Carroll RC, Grody M, Vronskaya S, Kandel ER. 2008. Transgenic mice lacking NMDAR-dependent LTD exhibit deficits in behavioral flexibility. Neuron 58:104–117.

Oliveira JF, Sardinha VM, Guerra-Gomes S, Araque A, Sousa N. 2015. Do stars govern our actions? Astrocyte involvement in rodent behavior. Trends in Neurosciences 38:535–549.

Panatier A, Robitaille R. 2016. Astrocytic mGluR5 and the tripartite synapse. Neuroscience 323:29–34.

Panatier A, Vallée J, Haber M, Murai KK, Lacaille J-C, Robitaille R. 2011. Astrocytes are endogenous regulators of basal transmission at central synapses. Cell 146:785–798.

Paxinos G, Franklin KBJ. 2001. The Mouse Brain in Stereotaxic Coordinates, Second Edition. 2 edition. San Diego: Academic Press.

Pinto-Duarte A, Roberts AJ, Ouyang K, Sejnowski TJ. 2019. Impairments in remote memory caused by the lack of Type 2 IP3 receptors. Glia 67:1976–1989.

Sardinha VM, Guerra-Gomes S, Caetano I, Tavares G, Martins M, Reis JS, Correia JS, Teixeira-Castro A, Pinto L, Sousa N, Oliveira JF. 2017. Astrocytic signaling supports hippocampal–prefrontal theta synchronization and cognitive function. Glia 65:1944– 1960.

Shrivastava AN, Kowalewski JM, Renner M, Bousset L, Koulakoff A, Melki R, Giaume C, Triller A. 2013. β-amyloid and ATP-induced diffusional trapping of astrocyte and neuronal metabotropic glutamate type-5 receptors. Glia 61:1673–1686.

Sun W, McConnell E, Pare J-F, Xu Q, Chen M, Peng W, Lovatt D, Han X, Smith Y, Nedergaard M. 2013. Glutamate-dependent neuroglial calcium signaling differs between young and adult brain. Science 339:197–200.

Umpierre AD, West PJ, White JA, Wilcox KS. 2019. Conditional Knock-out of mGluR5 from Astrocytes during Epilepsy Development Impairs High-Frequency Glutamate Uptake. J Neurosci 39:727–742.

Viana JF, Machado JL, Abreu DS, Veiga A, Barsanti S, Tavares G, Martins M, Sardinha VM, Guerra-Gomes S, Domingos C, Pauletti A, Wahis J, Liu C, Calì C, Henneberger C, Holt MG, Oliveira JF. 2023. Astrocyte structural heterogeneity in the mouse hippocampus. Glia 71:1667–1682.

Wang Q, Kong Y, Wu D-Y, Liu J-H, Jie W, You Q-L, Huang L, Hu J, Chu H-D, Gao F, Hu N-Y, Luo Z-C, Li X-W, Li S-J, Wu Z-F, Li Y-L, Yang J-M, Gao T-M. 2021. Impaired calcium signaling in astrocytes modulates autism spectrum disorder-like behaviors in mice. Nat Commun 12:3321.

Wang X, Lou N, Xu Q, Tian G-F, Peng WG, Han X, Kang J, Takano T, Nedergaard M. 2006. Astrocytic Ca2+ signaling evoked by sensory stimulation in vivo. Nat Neurosci 9:816– 823.

Xu J, Zhu Y, Contractor A, Heinemann SF. 2009. mGluR5 Has a Critical Role in Inhibitory Learning. J Neurosci 29:3676–3684.

Zhao Y-F, Ren W-J, Zhang Y, He J-R, Yin H-Y, Liao Y, Rubini P, Deussing JM, Verkhratsky A, Yuan Z-Q, Illes P, Tang Y. 2022. High, in Contrast to Low Levels of Acute Stress Induce Depressive-like Behavior by Involving Astrocytic, in Addition to Microglial P2X7 Receptors in the Rodent Hippocampus. International Journal of Molecular Sciences 23:1904.

Zhuang X, Oosting RS, Jones SR, Gainetdinov RR, Miller GW, Caron MG, Hen R. 2001. Hyperactivity and impaired response habituation in hyperdopaminergic mice. Proceedings of the National Academy of Sciences 98:1982–1987.

